# Evolution of opsin genes among butterfly species from different forest strata

**DOI:** 10.1101/2025.06.13.659549

**Authors:** Joséphine Ledamoisel, Andrew Dang, Julien Devilliers, Tiphaine Marvillet, Sophie Lemoine, André V. L. Freitas, Carolina Pardo-Diaz, Erika Páez, Manuela Lopez-Villavicencio, Adriana Briscoe, Vincent Debat, Violaine Llaurens

## Abstract

How do adaptive processes act on the coevolution of opsin genes shaping visual capacities in animals? At large phylogenetic scales, natural selection due to variations in light environments has been found to impact the evolution of photoreceptor genes. However, sexual selection and male-male competition may also influence their evolution. Here, we investigate the diversification of opsin sequences and their expression in closely-related blue *Morpho* butterfly species, living in different vertical strata within tropical forests, to shed light on the biotic, and abiotic selective pressures shaping the evolution of opsin genes. We combined comparative genomics, transcriptomics, and immunochemistry to characterise the expression and the spatial distribution of the opsin proteins found in the eyes of *Morpho helenor*. We detected a double duplication of the LW opsin and found unique ommatidial types compared to other butterfly species. We then investigated the evolution of opsin genes among 21 *Morpho* species, found signatures of positive selection on two opsin genes (LW3Rh and BRh), and identified key co-evolving amino-acids within and among these two opsin genes. We showed that opsin evolution was correlated with both light environment and wing coloration, highlighting the effect of several selective pressures in the coordinated evolution of these proteins.

## Introduction

The sensitivity to different wavelengths of light, linked to variations in opsin genes, has been extensively studied in multiple species, revealing striking diversification of colour discrimination capacities in animals. Yet, in most species, the selective pressures imposed by the light habitat and ecological interactions within and among species that shape opsin gene family diversification remain largely unknown. Studying the diversification of the number and sequences of opsin genes in closely-related species may shed light on the effect of adaptation to different ecological niches in the evolution of this gene family playing a key role in visual capacity.

Variations in the amino-acid sequences of opsins can directly influence the sensitivity to different light wavelengths of the opsin-chromophore complex known as a visual pigment or rhodopsin. The combination of opsins expressed within and among the different photoreceptor cells within the eye then enables colour discrimination [1]. In insects like flies and butterflies, the chromophore bound to the opsin protein is 11-*cis*-hydroxy retinal [2] and the evolution of the light-absorbing properties of each visual pigment is linked to the nature of amino acids located around this chromophore [1]. Evidence of concerted evolution, due to gene duplication, mutations and gene conversion has been reported in the opsin gene family [3]. Selection acting on visual capacities may indeed trigger joint evolution of several opsin genes.

In Lepidoptera, phylogenetic reconstructions inferred an ancestral visual system with three main opsin proteins [4]: ultraviolet-sensitive opsins (UVRh), blue-sensitive opsins (BRh), and long wavelength-sensitive opsins (LWRh). However, in many butterfly families, multiple events of opsin gene duplication and mutation affecting the chromophore-interacting amino-acids have been documented [5,6]. The duplication of opsin genes, followed by shifts in their spectral sensitivity, has been shown to result in a finer sampling of wavelengths within the UV-visible (300-700 nm) wavelength range, as for instance in Nymphalidae [7] or Lycaenidae [8]. The compound eyes of butterflies are composed of ommatidia, formed by a cluster of nine photoreceptor cells (named R1 to R9), which contain different sets of opsin proteins. Typically, R1 and R2 express either UV or blue opsins while R3-8 express long-wavelength opsins, although there are exceptions. Variations in the spatial expression of the different opsins within the different cell types, as well as in the spatial distribution of ommatidial types throughout the compound eyes, also strongly influence the evolution of sensitivity to light and colour discrimination [5]. The co-expression of opsins in R1 and R2 cells has been documented in *Heliconius* butterflies, with LWRh and BRh co-expression participating in spectral-tuning [9]. The evolution of colour discrimination might therefore stem from various molecular and cellular processes: reconstructing the evolutionary steps leading to divergent visual systems thus requires not only tracking opsin duplications and the joint evolution of their amino-acid sequences, but also investigating their levels of expression and their spatial distribution within and among ommatidia.

Identifying the respective contribution of the different selective pressures acting on the evolution of opsin spectral properties and spatial distribution within the eyes is challenging, because vision is involved in multiple fitness-related traits in many animals, such as foraging [10], mate recognition [11] and/or orientation [12]. Divergent evolution of opsins expression has been shown among species living in different light environments: for example, in aquatic environments, opsins were found to be differentially expressed among cichlids [13] and cardinalfish [14] species living in different photic micro-habitats (shallow *vs*. deep waters). Ecological interactions can also shape the evolution of colour discrimination capacities in animals. Visual cues are often used in mate choice and species recognition (in invertebrates [15] and in vertebrates [16]), potentially promoting the selection of different visual sensitivities in different taxa. As the display of bright colorations is often associated with traits contributing to fitness, sexual selection could favour the evolution of specific colour sensitivity ([17], [18]). The evolution of colour discrimination could also be promoted between closely-related species sharing the same habitat, as poor species recognition could lead to substantial deleterious reproductive interference [19], although the literature still lacks empirical evidence.

The rise of genomics has permitted the study of the evolution of opsin sequences at very large phylogenetic scales [20,21], highlighting an important effect of the light environment on the macro-evolution of visual systems [22]. However, sexual selection is more likely to influence opsin evolution at a smaller phylogenetic scale, as revealed in fishes [23] and butterflies [24]. Investigating the diversification of opsin gene sequences and patterns of expression in closely-related species with contrasted ecologies can thus shed light on the respective effects of biotic and abiotic selective pressures shaping the evolution of the opsin gene family.

Here, we focus on the evolution of opsin proteins within the neotropical butterfly genus *Morpho* (Nymphalidae), where sympatric species are distributed across different vertical forest strata [25], from the darker understory to the brighter canopy, as well as different temporal niches [26], from dawn specialized to noon-specialized species. Such specialisation into different spatio-temporal microhabitats is associated with a strong heterogeneity in the light environment [27]. The divergent light environment encountered by different *Morpho* species could thus influence the evolution of their colour discrimination capacities. Furthermore, these different species strongly differ in the structural and pigmentary colours of their wings. Behavioural experiments carried out in blue *Morpho* species showed that iridescent blue coloration can be a cue used by males during both intra- or interspecific interactions, like male-male competition and female recognition [26,28]. Consequently, as shades of iridescent blue vary between species [29], different sensitivities to blue light could have evolved within this genus. Indeed, some *Heliconius* species displaying UV-coloured pigments on their wings have accurate UV colour discrimination due to duplicated *UVRh* [30]. Visual modelling combined with behavioural experiments revealed that this additional violet receptor could facilitate the discrimination of conspecifics during mate choice [31,32], highlighting the effect of sexual interactions affecting opsin evolution in butterflies.

Most nymphalid butterfly species express three spectral classes of visual pigment in their eyes, encoded by single-copy *UVRh*, *BRh* and *LWRh* opsin genes. The recent sequencing of the genomes of three blue *Morpho* species has revealed duplications of the *LWRh* opsin gene resulting in three copies, in sharp contrast with other closely-related Satyrinae butterflies [33]. This result raises the question of the evolutionary origin of such duplications and of the selective regimes acting on the five opsin genes throughout the *Morpho* genus. Recent studies also investigated the different light sensitivities of the photoreceptor cells in the blue species *Morpho helenor* [34,35], calling for an investigation of the spatial distribution of opsins in different photoreceptors.

We thus investigate the evolution of opsin gene sequences and expression in the genus *Morpho* and aimed at detecting signature of different selective factors on the evolution and co-evolution of opsin genes. We first conducted a detailed study of the visual system of the only currently bred species *M. helenor* by characterizing all the opsin genes found in its genome, quantifying opsin mRNA expression in the eyes of males and females, and localizing the proteins corresponding to the UVRh, BRh and LW1Rh opsins within the photoreceptor cells in different ommatidia composing their retina. We then investigated the evolution of opsin genes among 21 *Morpho* species (out of the 30 described species) using samples captured in the wild, by (1) testing for signature of selection against the null hypothesis of neutral evolution and (2) assessing how the co-ordinated evolution of amino-acids within and among opsins could shape the diversification of their visual systems. In particular, we tested whether the evolution of their photosensitive proteins is correlated to their ecological environment or the evolution of their wing coloration. Our study thus aims at identifying the ecological and molecular mechanisms driving the diversification of visual systems.

## Materials and Methods

### Characterisation of opsins expression and spatial localisation in *Morpho helenor*

#### Testing RNA differential expression between males and females in M. helenor

We used the previously assembled genome of *M. helenor* [33] to retrieve the sequence of the 5 opsin genes (*BRh*, *UVRh*, *LW1Rh*, *LW2Rh* and *LW3Rh*) found in this species. In order to test whether opsin genes were expressed in *Morpho helenor* tissues, the eye transcriptomes of 5 males and 5 females *M. helenor theodorus* were sequenced. The specimens were purchased from a breeding farm located in Ecuador (Quinta De Goulaine, https://quintadegoulaine.com/es/papillons.php) and raised in insectaries at the Smithsonian Tropical Research Institute in Gamboa, Panama, between January and March 2023. The Qiagen RNAeasy Mini Kit was used to extract the RNA of each eye sample following the manufacturer’s instructions. The detailed protocol used for library preparation and Illumina sequencing is presented in the supplemental methods. The *M. helenor* genome annotation [33] was used to compute gene expression. All overlapping regions between alignments and referenced exons were counted and aggregated by gene using HTSeq-count 0.5.3 [36]. All counts with less than 10 reads were filtered out. The sample counts normalization (using the median of ratio method) and differential analyses to compare opsin expression between males and females were performed using DESeq2 1.42.1 [37]. The *p*-value was adjusted using the Benjamini-Hochberg method with a threshold set at 0.05.

#### Spatial distribution of opsin proteins within M. helenor eyes revealed by immunohistochemistry

We used an immunohistochemistry approach to locate the opsin proteins within the eyes of *M. helenor*. First, we used the translated amino acid sequences of *M. helenor* UVRh, BRh, and LWRh opsins to identify antigenic, accessible, and unique peptides as candidate targets for polyclonal antibody production. An antibody against the peptide GIVKQVFAHEAALRE in the loop domain between helix 5 and 6 of UVRh was generated in guinea pigs, an antibody against the peptide GWNIPEEHQDLVHE in the N-terminus domain of BRh was generated in goat, an antibody against a peptide GSDTGPGISC in the N-terminus domain of LW1Rh was generated in rabbit (Biosynth International, Gardner, MA, USA). Although we attempted to make a guinea pig antibody for LW2Rh using the peptide AAPDENG and a chicken antibody for LW3Rh using the peptide FDTSLQNIL, the immunopurified antibodies did not label eye tissue. We decided to focus our spatial expression analyses on BRh, UVRh and LW1Rh.

*M. helenor* pupae used for immunohistochemistry were purchased from Costa Rica Entomological Supply. Upon delivery, pupae were hung in greenhouse enclosures where they were housed post-eclosure and until processing. Butterflies freely flew in the enclosures and fed on moistened orange slices. Five male and five females were processed for immunohistochemistry. Methods adapted from previous studies [24,32,38,39] are presented in the supplemental method file. To validate the specificity of our custom antibodies, western blot analyses were also performed on protein extracts of *M. helenor* eyes following the protocol described in the supplemental method file by Boster Bio (Pleasanton, CA).

### Investigating the evolution of opsin sequences throughout the *Morpho* genus

#### Retrieving the opsin sequences from 20 Morpho reference genomes

The visual opsin sequences of 8 *Morpho* species observed in the understory habitat (*M. helenor*, *M. marcus*, *M. eugenia*, *M. granadensis*, *M. amathonte*, *M. menelaus*, *M. deidamia*, *M. achilles)* and 3 *Morpho* species found in the canopy habitat (*M. telemachus*, *M. hecuba, M. rhetenor)* were retrieved from the annotations of the *Morpho* genome assemblies [33,40]. Additional genomes including 4 understory species (*M. godartii, M. sulkowskyi, M. lympharis, M. epistrophus*) and 5 canopy species (*M. theseus*, *M. niepelti* and *M. cypris, M. cisseis, M. anaxibia*) were specifically sampled in the wild for this study and their sequencing, assembly and annotation were performed following the same pipeline as described in [40]. Among the 30 *Morpho* species described in the literature [25], we thus had access to the genomic data of 20 *Morpho* species scattered across the phylogeny. Overall, one *UVRh*, one *BRh* and three *LWRh* (referred to as *LW1Rh*, *LW2Rh* and *LW3Rh* hereafter) opsin genes were identified within each of these 20 genomes.

#### Amplification of additional opsin sequences throughout the Morpho genus

In order to retrieve the opsin sequences of the *Morpho* species for which assembled genomes are not available, the coding sequences of the *UVRh*, *BRh* and *LWRh* opsins from *M. helenor* and *M. telemachus* were used to design specific PCR primers (see Table S1) targeting these three genes with Primer3 [43]. PCRs were performed on 10 *Morpho* additional species (*M. amphitryon*, *M. hercules*, *M. aurora*, *M. absoloni*, *M. rhodopteron*, *M. portis*, *M. aega*, *M. zephyritis*, *M. iphitus*, *M. polyphemus*) using DNA samples from collection specimens stored in the National Museum of Natural History in Paris, France. The DNA of those *Morphos* was extracted using the Qiagen extraction DNeasy Blood Tissue Kits. Details on the used PCR solution and thermal cycling conditions can be found in the supplemental methods.

The PCR products were sequenced using the short-read Illumina technology. The Geneious Prime program (2022.1.1 version) was then used to reconstruct the whole opsin sequences. Using this method, we retrieved partial *LWRh* opsin sequences from one new understory species (*M. aurora*), adding to the 20 *Morpho* species already studied from *Morpho* genomes.

#### Sequencing of eye transcriptome of 6 Morpho species

To check for opsin RNA expression in different *Morpho* species scattered across the phylogeny of the genus, the eyes of freshly caught males belonging to 5 sympatric species from French Guiana (*M. achilles*, *M. deidamia*, *M. hecuba*, *M. rhetenor* and *M. marcus*), were dissected and stored in RNAlater at −80°C before extraction. RNA was extracted using the Qiagen RNeasy Mini Kit according to the manufacturer’s instructions and stored at −80 °C until use. The transcriptome of each sample was sequenced using Nanopore technology Library preparation and the Nanopore sequencing protocols are presented in the supplementary methods. The extracted reads were assembled on their respective previously-assembled and annotated reference genomes [33,40] and the RNA counts were extracted using HTSeq-count 0.5.3 [36]. As we collected data for only one individual per species, this analysis was used to assess the presence of expressed opsin genes in the eyes of *Morpho* but not their absence, using TPM counts.

#### Sequence alignment and tree reconstruction

The coding opsin sequences of each opsin, retrieved from 21 *Morpho* species, were aligned using the MUSCLE algorithm [41] implemented in MEGA X [42]. IQtree [43] was used to generate the *UVRh*, *BRh*, *LW1Rh*, *LW2Rh* and *LW3Rh* phylogenies, as well as a *LWRh* tree reuniting the 3 long-wavelength copies. The ModelFinder option [44] was used to select the best respective evolutionary models for each tree. Nodes’ robustness was estimated using Ultrafast Bootstrap [45] on 10000 iterations.

#### Detection of signature of selection acting on the evolution of opsin sequences

To characterise the selection regime acting on the evolution of opsin sequences and assess departure from neutral evolution, we studied the ratio between the non-synonymous (*dN*) and synonymous (*dS*) mutations in the coding sequences of the different opsins ⍵*=dN/dS*. A gene with a higher non-synonymous substitution rate compared to its synonymous substitution rate (⍵>1) is considered positively selected. The *codeml* program from the software package PAML 4.9j [46] was used to test for gene-wide pervasive selection acting on the evolution of visual opsins. To investigate whether selection is acting on specific codons in every branch of the opsin gene trees, we performed model comparisons using models proposed by PAML (Site Models M0, M1a, M2a, M7, M8). In particular, the M2a and M8 models were used to assess positive selection on each opsin gene, by comparing their prediction to their respective null models M1a and M7. A Bayes Empirical Bayes (BEB) analysis was used to identify the positively selected amino acids among the genes under positive selection. As PAML’s Site Model is not suited to detect gene-wide selection occurring only in a few subsets of branches and sites, we also used the BUSTED program [47] from the software HYPHY v2.5.65 [48] to test for episodic selection on *Morpho* opsin genes. This method can detect signatures of selection happening on at least one branch. The gene trees used for PAML’s site model and BUSTED are available in Figure S1. Additionally, we used a clade model (PAML’s Cmc Model: M2a_rell (null) vs. Model C) on the *LWRh* duplication tree to test for a signal of differential evolution between the 3 *LWRh* opsin copies using a three-partition model.

Finally, in order to test the effects of ecological factors on the evolution of opsin sequences, we used a clade model (PAML’s Cmc Model) to analyse putative differential evolution of the opsin genes between species sharing an iridescent phenotype and those that do not (Iridescent vs. Non-Iridescent), or between species living in the canopy or the understory (Canopy vs. Understory). When a signal of differential evolution was found, we used HYPHY’s RELAX model [49] to test whether selection has relaxed or intensified in the selected branches.

#### Detection of correlated amino acid evolution

We used the Evo-Scope pipeline [50] to determine whether the evolution of some amino acids in a given opsin protein could (i) influence the evolution of other amino acids within the same protein or (ii) impact the evolution of the sequence of other opsin proteins. The Evo-Scope pipeline was designed to study correlated evolution of any discrete trait by accounting for the phylogenetic structure of the data. Applying this method to amino-acid changes allows to identify the correlations in amino acids changes during the diversification of *Morpho* species, without any *a priori* on the physiochemical constraints preventing certain combinations to persist.

In order to apply this method to our data, we considered each amino acid of the LWRh, BRh and UVRh opsin proteins as a discrete trait to analyse the co-occurrence of amino acid shifts found across the species phylogeny. *P-*values assessing the probability for an amino acid pair to be under correlated evolution were adjusted using the Benjamin-Hochberg (BH) to control the false discovery rate (FDR). We then retained amino acid pairs exhibiting a high signal of correlated evolution (*p-value* < 0.0001). A permutation test with 100 000 iterations was performed to assess whether pairs containing positively selected amino acids were more likely to be found than random combinations. Statistical analyses were run in R version 4.3.0.

#### Inferring putative functional roles to the amino acid changes observed in the genus Morpho

First, we identified which amino acid changes can be associated with changes in spectral sensitivity, by cataloguing tuning sites within the *BRh* and *LWRh* opsin sequences using the current literature on functional effects of opsin amino-acid variations in other butterflies [7,51–54]. We then used the MUSCLE algorithm [41] implemented in MEGA X [42] to align opsin sequences of *Morpho* and other butterfly species to locate the position of candidate tuning-sites. This method allows us to assess whether the amino acid detected as under positive selection correspond to candidate tuning sites putatively affecting sensitivity to different wavelengths.

We then located the positively selected amino acid sites and the sites associated with a signal of correlated evolution within the protein and specifically their proximity to the chromophore. To this aim, we predicted the 3D structure of the different opsins of *M. helenor.* The opsin amino acid sequences were uploaded to the SWISS-MODEL protein recognition engine [55] to generate a template aligned against the invertebrate jumping spider rhodopsin crystal structure [56]. The protein structures were edited in PyMOL (The PyMOL Molecular Graphic System, Version 2.6 Schrödinger, LLC) and the sites within a range of 5A of the 11-*cis*-retinal binding pocket were visualized via the predicted homology model.

## Results

### Opsin diversity, expression and localisation in the eye of M. helenor

The analysis of the Illumina short read RNA-seq data of the eye tissues of *M. helenor* revealed that all five opsin genes (*UVRh*, *BRh*, *LW1Rh*, *LW2Rh* and *LW3Rh*) are expressed in the eye tissues of both males and females (Table S2, Figure S2.A), without differential expression detected sexes (Table S3 and Figure S2.B).

When analysing opsin protein expression, two of the five custom polyclonal antibodies (anti-LW2Rh and anti-LW3Rh) generated against the opsins did not yield any staining when tested on eyes. Using the remaining three antibodies that did work (see Figure S3 for western blot results), we characterized the location of the UVRh, BRh and LW1Rh opsin proteins within *M. helenor* eyes using immunochemistry. We found UVRh, BRh, and LW1Rh opsin proteins to be expressed in the R1 and R2 photoreceptors of both female and male specimens of *M. helenor* (Figure 1 A & B). No opsin expression was detected in the R3-8 cells with these three antibodies While co-expression of BRh and LWRh opsins in some R1 and R2 cells has been reported along a dorsal-ventral gradient in the *Heliconius melpomene* eye, with co-expression peaking in the ventral retina and expression of LWRh-only in some R1 and R2 cells dorsally [57], we found no evidence of co-localization of BRh and LW1Rh in the part of the *M. helenor* retinas examined here.

**Figure 1.**
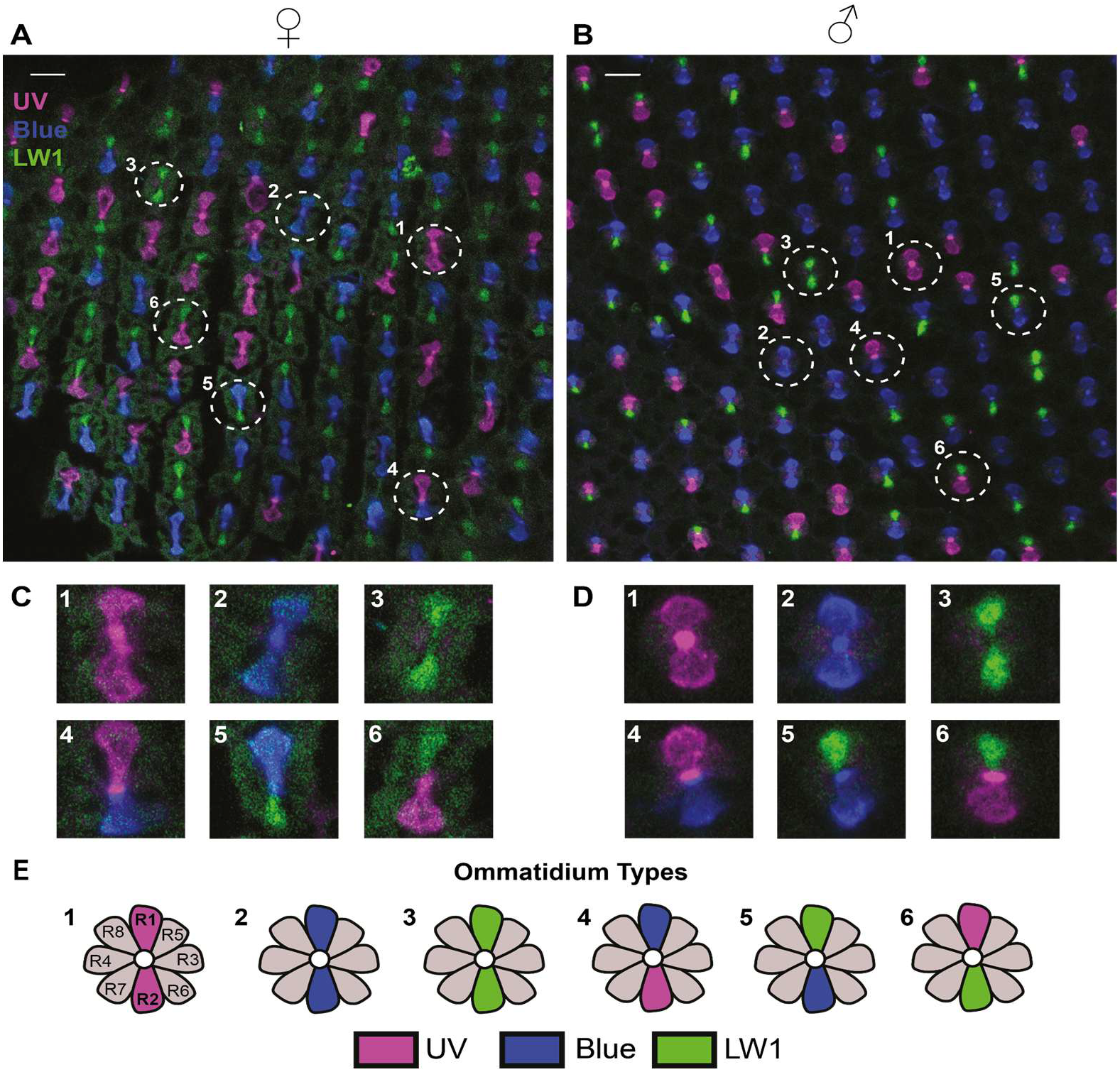
Characterisation of opsin protein expression in the R1 and R2 cells of different ommatidia of male and female *M. helenor* eyes. Anti-UVRh (magenta), anti-BRh (blue), anti-LW1Rh (green), antibody staining of (A) female and (B) male *M. helenor* retinas. Scale bars represent 20 microns. Individual ommatidium types identified in (C) female and (D) male retinas based on BRh, LW1Rh, and UVRh opsin expression. (E) Cartoon of the at least six ommatidia types based on opsin expression in the R1 and R2 photoreceptor cells of *M. helenor*.

Sex differences in ommatidia type and photoreceptor type distribution were tested with Poisson GLMMs with individual animals as a random effect (Figure S4). Raw cell counts for each individual and cell type are provided in Table S4. The distribution of ommatidia types differed significantly between males and females (sex × type interaction, likelihood ratio test: χ² = 27.5, df = 5, *p-value* < 0.001). Pairwise comparisons revealed a significantly higher proportion of LW1Rh-LW1Rh ommatidia in females (adjusted *p-value* = 0.015), whereas BRh-BRh ommatidia were more frequent in males (adjusted *p-value* = 0.041). In a similar trend, the distribution of photoreceptor types is also significantly different between sexes (χ² = 17.7, df = 2, *p-value* < 0.001). LW1 photoreceptors were significantly more frequent in females (adjusted *p-value* = 0.009), whereas blue photoreceptors were more frequent in males (adjusted *p-value* = 0.022).

Overall, our results revealed a sexual dimorphism in the proportion of photoreceptor and ommatidia types. Since our analysis only focused on BRh, UVRh, and LW1Rh, an even more complex retinal organisation is expected, but additional analyses would be needed to confirm it.

### Opsin diversification at the genus scale in Morpho butterflies

After characterising the spatial expression of opsin gene within the only species of *Morpho* that can be bred in captivity, we then explored the variations in the opsin sequences in other *Morpho* species, using samples collected in the wild. The *UVRh*, *BRh* and *LWRh* genes detected in the genome of *M. helenor* were also found in the assembled genomes of 20 other *Morpho* species. In addition, partial LW2Rh and LW3Rh sequences were also retrieved for *M. aurora* using PCR (Figure 2). The two duplication events in the *LWRh* gene are thus shared throughout the genus, but are not found in a closely-related Satyrinae outgroup species (*Elymnias hypermnestra*), implying these *LWRh* duplications happened shortly before the diversification of the *Morpho* genus. The analysis of the transcriptome of the eyes of *M. achilles, M. deidamia, M. hecuba, M. rhetenor* and *M. marcus* males showed that all five opsin genes are expressed in the eyes of those species (Table S5), suggesting that these five opsin genes could be involved in colour vision in understory species (*M. achilles, M. deidamia*), canopy species (*M. hecuba, M. rhetenor*) and phylogenetically basal *Morpho* species (*M. marcus*). The phylogenetic reconstruction of the *LWRh* genes of *Morpho* butterflies (Figure S5) then showed that *LW1Rh* and *LW2Rh* are more closely-related to each other than they are to *LW3Rh*.

**Figure 2:**
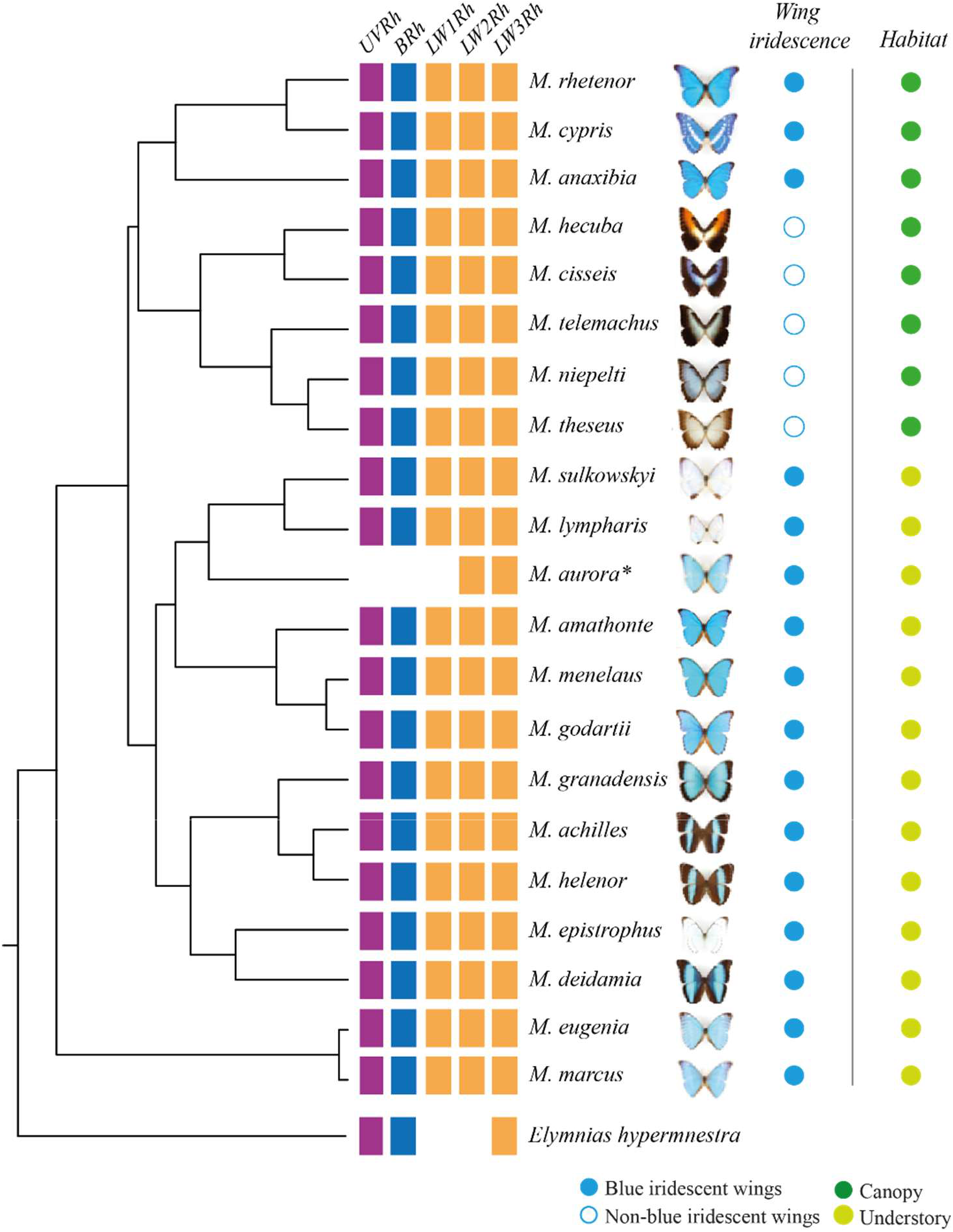
Distributions of opsin duplications and traits (wing coloration and microhabitat) in the 21 species of the genus *Morpho* used in this study. The phylogenetic relationships among the studied species were retrieved from [25]. The coloured tiles show the *UVRh*, *BRh*, *LW1Rh*, *LW2Rh* and *LW3Rh* opsin genes for each *Morpho* species. The opsin sequences retrieved from the species marked with black asterisks were amplified using PCR and are thus only partial sequences. The male dorsal wing pattern (Wing colour: blue circles = blue-iridescent wings *vs.* white circles = non-blue-iridescent wings), and the habitat (Habitat: light-green circles = understory species *vs.* dark-green circles = canopy species) is provided for each studied species. The opsin genes of the outgroup *Elymnias hypermnestra* are also pictured (GenBank GCA_015832385.1).

### Variations at spectral tuning sites: evidence of putative spectral sensitivity shifts in BRh and LWRh rhodopsins in the Morpho genus

Because the amino acid sequence of a photoreceptive protein determines its capacity to absorb specific wavelengths, we checked whether variations of amino acids observed in *Morpho* species were associated with tuning sites known in other butterflies in BRh and LWRh opsins. In the rest of this article, amino acid positions are numbered according to the *Morpho* sequences. An equivalence table with the numberings from the original sources [7,51–54], and the putative effect of amino acid changes, is provided in Table S6.

Briefly, *Morpho* BRh opsin sequences show amino acid substitutions at sites associated with blue shifts in butterfly blue rhodopsin absorbance spectra such as S135A [52,54], T301A, F195Y, and K216 [51]. Five species (*M. anaxibia, M. amathonte, M. godartii*, *M. rhetenor* and *M. cypris*) share a putative blue-shifting F195Y substitution (see Figure S6 and Figure S7). Interestingly, a further T301A predicted blue-shifting substitution [51] is only found among *M. rhetenor* and *M. cypris*, suggesting that some amino acid shifts are unique to blue canopy species.

In the LWRh opsins, a S185A putative blue-shifting amino acid substitution [7] (equivalent to S180A in human M/L opsins [58]) occurred in the LW1Rh and LW2Rh lineages, but not in the LW3Rh opsin, suggesting that LW3Rh may retain sensitivity to longer wavelengths. Interestingly, a A185S reversion was found in the LW1Rh opsin of the non-blue canopy species *M. hecuba* and *M. cisseis*, which could thus have a LW1Rh protein with a closer sensitivity to the LW3Rh protein compared to other *Morpho* species. Among LW3Rh, an A137G potentially red-shifting substitution (equivalent to site 116 in bovine rhodopsin) [53] is also found in five canopy and understory species (see Figure S5 and Figure S8). Overall, these results suggest that the duplicated *Morpho* LWRh rhodopsins (especially LWRh1/2 vs. LWRh3) might be sensitive to different wavelengths, consistent with a diversifying evolution of their visual system.

### Signature of positive selection on the BRh and LW3Rh Morpho opsin genes

We then studied the ratio between non-synonymous and synonymous substitutions in the coding sequence of the different *Morpho* opsins (*dN/dS*=*ω*) to characterize the selection regime acting on the evolution of the five opsin sequences. A signal of pervasive positive selection was found on the *BRh* (*ω_BRh_* = 2.21, *p-value* = 0.0003) and *LW3Rh* (*ω_LW3Rh_* = 4.48, *p-value* = 0.0001) opsin genes using site models M8 implemented in PAML (Table S7). We also found significant evidence of episodic selection on *LW3Rh* (*p-value* = 0.009) using BUSTED at a gene-wide scale (Table S8). PAML’s BEB was run on those two genes to identify specific positively selected amino acid sites at a 0.75 probability cut-off (Table S7); the sites 85, 113, 189, 232, 246, 363 in BRh, and 88, 129, 136, 180, 227, 229, 236, 305 in LW3Rh are found to be under positive selection. While none of these sites corresponds to known tuning sites in other butterfly species, they are either located near those sites or in the vicinity of amino acids identified with homology modelling as part of the binding pocket of the chromophore (see Figure S7, S8 and S9). In LW3Rh in particular, the amino acid 227 is part of the binding pocket of the protein.

To investigate the potential change in evolutionary rates among the three *LWRh* opsin duplications, we performed a clade model and found evidence of differential evolution between duplications (*ωLW1=0.32, ωLW2=0.16, ωLW3=0.57, p-value < 0.0001,* Table S9).

### Testing for ecological factors influencing the evolution of Morpho opsins

We looked for shifts in selective regimes acting on the sequences of the five *Morpho* opsin genes depending on the species ecologies, using the CmC PAML model. We found a significant statistical effect of wing coloration and habitat on the evolution of the *LW2Rh* and *LW3Rh* genes (Figure 3, statistical analyses in Table S10), with the species exhibiting iridescent wings having a higher *ω* than the non-iridescent species (*LW2Rh*: *ω_LW2Rh_iridescent_* = 0.65 *vs. ω_LW2Rh_Non-iridescent_* = 0.14 and *LW3Rh*: *ω_LW3Rh_iridescent_* = 0.95 *vs*. *ω_LW3Rh_Non-iridescent_* = 0.07), and with the species living in the understory having a higher *ω* than the species living in the canopy (*LW2Rh*: *ω_LW2Rh_understory_* = 1.13 *vs. ω_LW2Rh_canopy_* = 0.00 and *LW3Rh*: *ω_LW3Rh_understory_* = 0.39 *vs*. *ω_LW3Rh_canopy_* = 0.00). We then tested whether those results could be due to relaxed or intensified selection using RELAX (Table S11), and found a significant signal of selection relaxation in LW3Rh for iridescent species (K=0.63, *p*-value = 0.034), suggesting that selection on this gene may be weakened in iridescent species.

**Figure 3:**
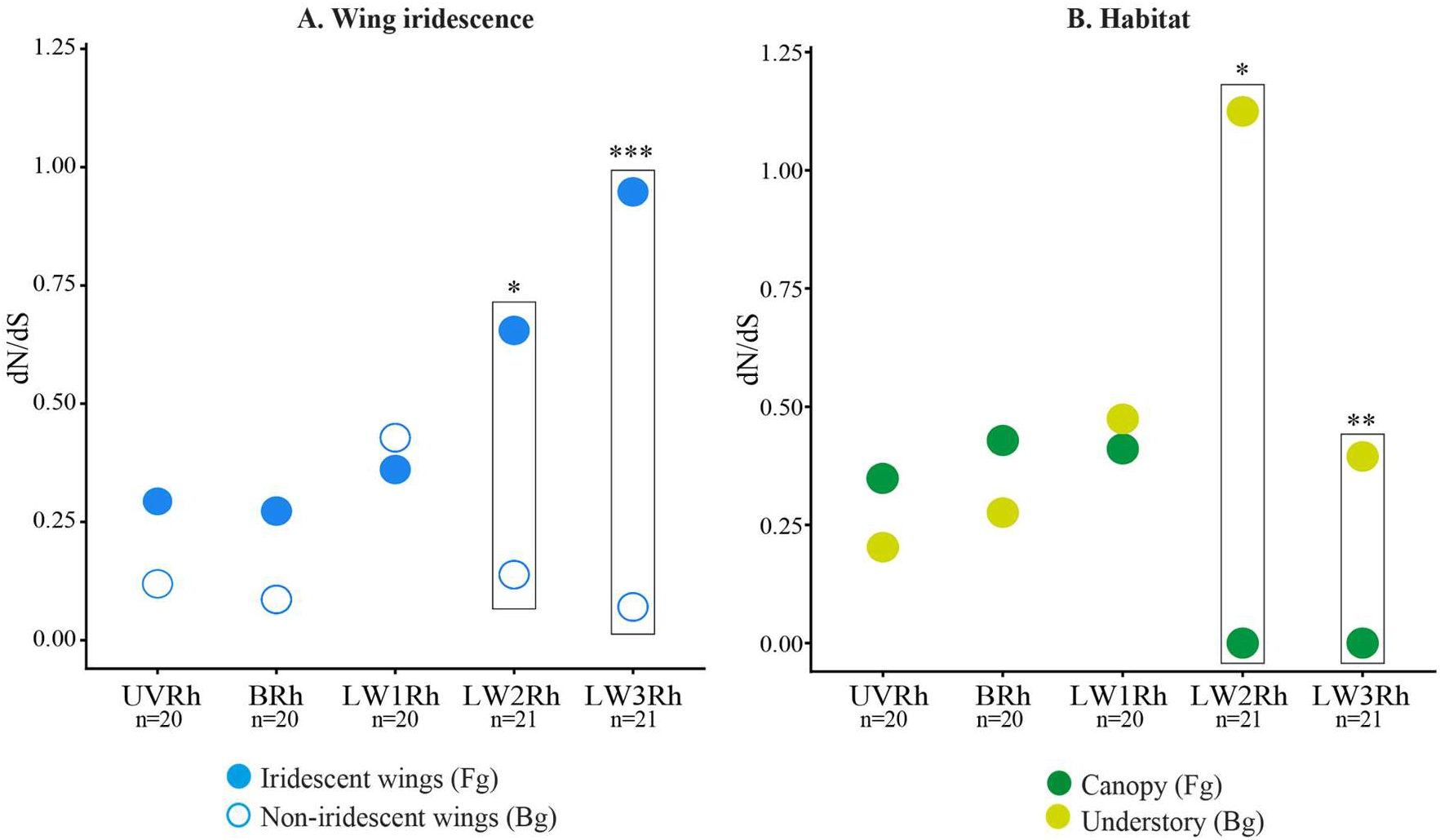
Variation of *Morpho opsin* gene *dN/dS* depending on wing coloration. **(A) and habitat (B).** The *dN/dS* was estimated by the CmC PAML model. The dN/dS of blue species are shown in blue, in white for non-blue species, in dark-green in canopy species, in light-green in understory (Fg = foreground, Bg = background). The number of analysed genes are shown below each opsin type.

### Evidence of correlated amino acid evolution between LWRh opsin genes

Finally, we used the Evo-Scope pipeline [50] to test for signatures of co-evolution of amino-acids within and among all five opsin proteins throughout the 20 species of the genus *Morpho* studied here.

Out of more than 3 million of possible pairs of amino acids, only 266 amino acid pairs showed a signature of correlated evolution (p-value<0.0001), among which 31.20% are located within the same protein, and 68.80% among different opsins (statistical analyses in Table S12). The top 5% most significant correlations (Figure 4) were found to be more enriched in positively selected amino acids compared to the random expectation (*p-value=0.003*, Figure S10). Among LW3Rh, the evolution of site 227 (part of the binding pocket and under positive selection) is found correlated to the evolution of site 224. In addition, the evolution of site 136 (under positive selection) is correlated to the evolution of site 137 (equivalent to site 116 in bovine rhodopsin, Table S6) identified as an important tuning site in butterflies [53]. Interestingly, coevolution between positively selected sites are not limited to the same protein but can also occur between opsin proteins, especially between LW3Rh and BRh; positively selected sites 305 (in LW3Rh) and 85 (in BRh) show one of the strongest signatures of correlated evolution. The evolution of site 217 in LW3Rh is also correlated to the evolution of the positively selected site 246 located on BRh.

**Figure 4:**
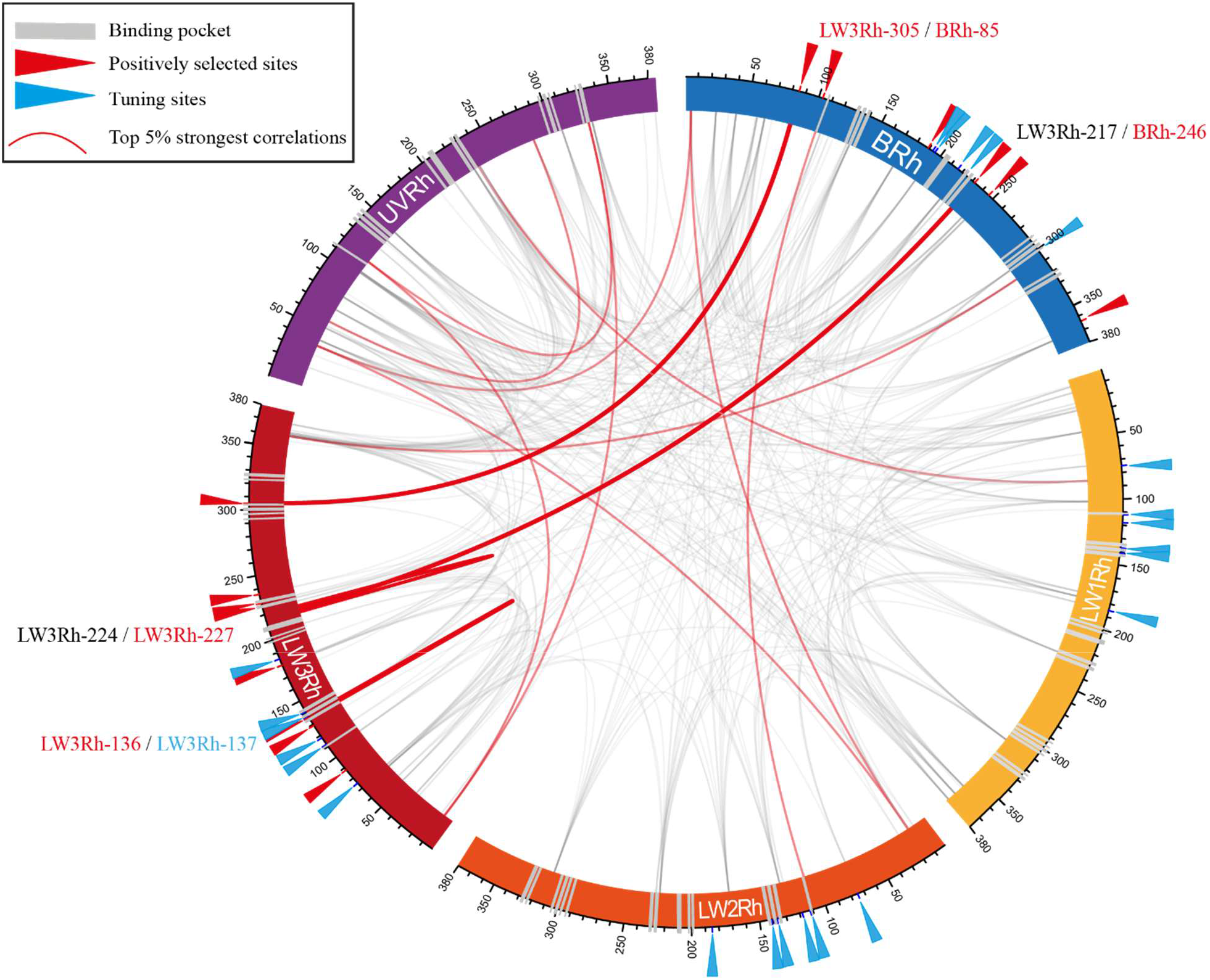
Correlated amino-acid evolution within and between the five opsin proteins observed throughout the genus *Morpho*. Circular mapping linking amino acid pairs presenting highly significant signatures of correlated evolution (*p-value*<0.0001). The tuning sites documented in other Lepidoptera and the positively selected sites detected in *dN/dS*-based analyses carried in Morphos are indicated on each protein using blue and red arrows respectively. Amino-acid correlations detected by EvoScope with *p-values* < 0.0001 are represented by grey arcs; the top 5% most significant *p-values* are linked by red arcs, among which the correlations involving positively selected sites are shown in bold.

Tuning sites documented in other butterflies, as well as sites detected as under positive selection were thus more likely to display strong signals of correlated evolution, consistent with a coordinated evolution of these amino acids, either due to compensatory mutations or joint evolution of opsin proteins enabling access to new wavelength sensitivity ranges.

## Discussion

### Diversification of expression patterns in duplicated opsins

The RNA expression analyses performed on *M. helenor* eye tissue confirmed that all 5 opsin genes, including the three different *LWRh* copies, are expressed in the eyes of *M. helenor* butterflies from both sexes. Our immunohistochemical assays then showed that UVRh and BRh opsins are expressed in R1 and R2 photoreceptive cells, consistent with previous observations in other butterfly visual systems (*e.g. Vanessa cardui* [59]; *Heliconius* [32]; *Danaus plexippus* [60]; *Pieris rapae* [61]). Most nymphalid butterflies express a single-copy *LWRh* opsin gene in their R3-R8 cells, with some nymphalids also expressing LWRh in some R1-2 cells [4]. Interestingly, expression of LW1Rh in *M. helenor* was restricted to the R1-R2 photoreceptor cells. This pattern of expression is consistent with the previous observation that LWRh is expressed exclusively in the R1-R2 photoreceptors of a subset of ommatidia in the dorsal eye of *H. melpomene* [57], and also suggests that duplicated *M. helenor LW1Rh* has a sub-functionalised domain of expression compared to the broader expression (R1-8 cell) of the ancestral single-copy *LWRh* gene. We did not observe any obvious co-expression of *UVRh, BRh* or *LW1Rh* in this part of the retina. While this is the first report of a duplicated *LWRh* gene specifically expressed in R1 and R2 cells in nymphalid butterflies, our observations are consistent with prior electrophysiological results. Polarization sensitivity measurements performed on the eyes of *M. helenor* [35], showed that R1 and R2 cells are strongly sensitive to UV and blue light, and are also sensitive to green or green/yellow light, which is likely mediated by LW1Rh. Although we were not able to characterize the pattern of expression of the LW2Rh and LW3Rh proteins in the eyes of *Morpho*, we assume these may be expressed in cells R3 to R8 because these cells are also sensitive the green and yellow light according to the polarisation sensitivity data [35]. We thus describe 6 different ommatidial types in *M*. *helenor* (Figure 1), but unknown LW2Rh or LW3Rh expression patterns will likely increase this number.

This peculiar expression pattern of opsins is also associated with a sexually dimorphic expression pattern of some opsins in *M. helenor*. In particular, higher LW1Rh photoreceptor count was found in the eyes of females. Furthermore, a high number of BRh photoreceptors was found in males *M. helenor* eyes, associated with a higher frequency of BRh-BRh ommatidia types in males compared to females (see Figure S4). Such sexual dimorphism in opsin protein expression has been reported in several taxa, with variation in opsin co-expression in different eye photoreceptor cells [62], a lack of opsin expression in a photoreceptor cell of one sex compared to the other [32], or variations in the number of photoreceptor cells [63]. Such sexual dimorphism could arise from different selective pressures affecting the evolution of the visual systems of males and females. For instance, the behaviours of *Morpho* males and females strongly differ: in the wild, males display a typical patrolling behaviour and show a striking response to blue stimuli [26]. On the other hand, females tend to fly in dense forest areas and to spend time locating and flying around host-plants. Those different ecological conditions encountered by males and females might generate divergent selection on visual systems. Long-distance detection of blue colour might be advantageous in males searching for females and might promote increased sensitivity to blue wavelengths. Although *M. helenor* males do not have duplicated *BRh* opsin genes as other blue sensitive butterflies (Lycaenidae [62]; Pieridae [54]), their retina is composed of a higher proportion of BRh-BRh photoreceptors than other blue butterflies [62], specifically in males. In contrast, discrimination in the green wavelength might be promoted in females, because of the significance of host-plant detection in this sex, as suggested in other butterflies with sexually-dimorphic visual sensitivities [31].

The novelty of the *M. helenor* eye organization (currently based only on the study of BRh, UVRh and LW1Rh) compared to other butterflies and the number of expressed opsin genes in its visual system shows a divergent evolution of opsin gene expression in *Morpho* butterflies as compared to other Nymphalidae, that may result in the evolution of colour discrimination capacities.

### Ecological drivers of visual gene evolution in Morpho

The five opsin genes described in *M. helenor* were also found in the genome of 19 additional *Morpho* species. RNA expression data confirmed that those five genes are expressed in the eyes of five tested species (*M. achilles*, *M. deidamia*, *M. hecuba*, *M. rhetenor* and *M. marcus*), suggesting that this specific visual system might be ancestral to the diversification of *Morpho*. Models testing for pervasive positive selection on the 5 opsin genes revealed significant signals of positive selection mostly on the sequences of *BRh* and *LW3Rh* genes. Since several positively selected sites detected in our study are located close to previously-documented tuning sites (see Table S6) and/or close to the binding pocket, our results are consistent with an effect of selection on the diversification of visual sensitivities in those genes.

First, the analysis of the amino acid sequence of the *BRh* gene in *Morpho* showed that five species exhibit amino acid changes involving known tuning sites (*M. anaxibia, M. amathonte, M. godartii*, *M. rhetenor* and *M. cypris*). Overall, we found that the evolutionary rate of the *BRh* opsin was not significantly different between canopy and understory species, and surprisingly did not find a correlation between the evolution of the *BRh* opsin sequences and the presence or absence of iridescent blue coloration on the wings of *Morpho* butterflies. Nevertheless, iridescence quantification showed that the peak of reflectance of *M. helenor* wings can vary from 350 nm to 500 nm (UV/purple to green) depending on the angle of illumination [28], while spectral sensitivity data suggest that the blue-absorbing cells of *M. helenor* might be sensitive to approximately 450 nm [35]. The iridescent blue reflected by *Morpho* wings might thus rather be processed by UV and long-wavelength-absorbing rhodopsins during mate searching.

In line with this hypothesis, we found that *LWRh* genes are very diversified among *Morpho*. In particular LW1Rh and LW2Rh tend to bear blue-shifting amino acids as compared to LW3Rh (Table S6). Although functional validation is needed, the LW3Rh protein might thus be responsible for the spectral sensitivity peak found in *Morpho* at 570 nm, while LW1Rh and perhaps LW2Rh are likely responsible for the spectral sensitivity peak found at 505 nm according to the Pirih et al. (2022) study. Long-wavelength sensitive opsin diversification has been reported in a number of taxa (Hemiptera [64]; Coleoptera [65]; Anurans [66]) including butterflies (*Papilio Xuthus* [4]), and is often associated with change in light habitats and/or niche differentiation [67]. In *Morpho*, we found that the *LW2Rh* and *LW3Rh* opsin genes of iridescent species have significantly higher *dN/dS* ratios than those of non-iridescent species, suggesting accelerated evolution of *LWRh* genes in iridescent blue species. Interestingly, we also found that the evolutionary rate of the *LW2Rh* and *LW3Rh* gene is significantly higher in the *Morpho* species living in the understory compared to those living in the canopy. The understory habitat is dominated by dense vegetation: as the discrimination of subtle long-wavelength colour variation might be advantageous in a forest environment in terms of foraging and oviposition [68], this habitat is likely an important driver of green/red colour discrimination in *Morpho*.

### Adaptive co-evolution of opsin proteins

Because wavelength discrimination by the visual systems depends on the shape of sensitivity peaks and on the distances among them, controlled by the respective absorbances of each rhodopsin, its evolution likely involves integrated changes across the different opsin genes. To assess this coevolution among proteins involved in colour discrimination, we specifically tested for correlated shifts in amino acids both within and among opsins. We specifically considered amino acids as discrete traits, similarly to other known methods [69].

Within each of the opsin proteins, numerous pairs of amino acid sites displayed a significant signal of correlated evolution (Table S12). Amino acid substitutions at a site could indeed compensate for deleterious substitutions at other sites of the protein [70] and amino acid changes could also be driven by adaptation. Amino acid variations in *LW3Rh* revealed correlated evolution between pairs of sites documented as tuning sites in butterflies (site 137 in *Morpho*, equivalent to site 116 in bovine rhodopsin), located in the binding pocket (site 227 in *Morpho*), and detected under positive selection in our independent *dN/dS* analyses carried out on *Morpho* gene sequences (sites 136 and 227 in *Morpho*). Overall, these results are consistent with coordinated adaptive evolution across multiple sites in this protein, likely contributing to modulation of wavelength sensitivity.

Moreover, we also found co-occurring mutations between amino acid sites belonging to different opsin proteins, suggesting concerted evolution among the five genes part of the *Morpho* visual system. Correlated mutations between proteins have been extensively observed between proteins part of a network or physically interacting with one another [71], but correlations in the evolution of genes sharing functional relationships are much less documented. Yet, visual perception can be seen as the result of the integration of visual signals absorbed by different rhodopsins, potentially creating an indirect link between the evolution of those genes. The fact that both positively selected sites (305 in LW3Rh and 85 in BRh) show a strong signal of correlated evolution implies that selection may impact the functional evolution of groups of genes involved in vision. For example, following an opsin duplication, the gain of function of the new protein could impact the selection regime of other opsin genes to maintain fine-tuned spectral sensitivity. In Lycaenidae, coordinated shifts of amino acids have indeed been found in the *BRh* and *LWRh* opsins, enhancing their visual perception [52]. Although our approach is only correlative and functional validation is needed, it underlines the significance of studying the molecular co-evolution of opsins, because of the extensive evolutionary tinkering involved in the diversification of visual systems.

## Data availability

Newly assembled genomes (PRJNA1481994), RNA-seq data (PRJNA1482776) and isolated opsin sequences (PRJNA1482348) will all be available on the NCBI database upon publication. Alignments and scripts used for this study are available on a Zenodo repository: https://zenodo.org/records/21337715?preview=1&token=eyJhbGciOiJIUzUxMiJ9.eyJpZCI6IjllMmI5ZThjLTQ5NzAtNDI3Yy1hMTY1LTJhY2ZjNmE4MWNkNSIsImRhdGEiOnt9LCJyYW5kb20iOiJmMDliYmUyZjgxY2E0Zjk0Y2ZlYjlmYTBhYzc2OGY1MyJ9.p6ZHcnpI9mUWnXvjtmZHcclaEZ3bAfnkMyv_3gupfIybytRu0AABBP7RT4Tv9NWqG-nNN5ZpqaoXwNY2nXSZkg (temporary link for reviewers).

## Acknowledgements

The authors would like to thank Guillaume Achaz and Maxime Godfroid for the advice provided on the use of the EvoScope pipeline. We are also grateful to Owen McMillan from the Smithsonian Tropical Research Institute (Panama) for providing facilities to raise *M. helenor theodorus* and to Lisa Mesrop for eye dissections. We thank Etienne Delannoy from the POPS facility (IPS2) for conducting with GenomiqueENS the adaptation of the ONT RNA-seq protocol. We thank Renato R. Ramos for helping with field work in Jureia. We thank Ludvine Brajon for her advice in bioinformatics. All the bioinformatic analyses were performed on the MeSU platform at Sorbonne-Université. Collect and access to genetic resources for Peruvian samples (*M. cisseis*, *M. theseus*, *M. godartii*, *M. sulkowskyi*, *M. lympharis*) were authorized by the SERFOR permit RD-000158-2024-DGGSPFFS-DGSPFS, for Ecuadorian samples (*M. niepelti*) by the INABIO permit MAATE-DBI-CM-2023-0298, for Colombian samples (*M. cypris*) by the permit No. 530 issued by the Autoridad Nacional de Licencias Ambientales (ANLA), and for Brazilian samples (*M. anaxibia*, *M. epistrophus*) by the ICMBIO collecting permits (10438-4 to 10438-9).

We exported the eyes of *M. helenor theodorus* from Panama using exportation permit number PA-01-ARB-028-2023. The access to genetic and transcriptomic resources from wild samples from French guiana were authorized under NOR: TECL2520938S / 964. Pupae of *M. helenor* were exported from Costa Rica under USDA permit numbers P526P-21-06503 and 526-25-157-05736. Brazilian specimens and the present study are registered at the Brazilian SISGEN (A24FA6E).

