## Supplemental Methods for "Evolution of opsin genes among butterfly species from different forest strata"

RNA opsin library preparation and Illumina sequencing methods

In order to test whether opsin genes were expressed in *Morpho helenor* tissues, the eye transcriptomes of five males and five females *M. helenor theodorus* were sequenced. The Qiagen RNAeasy Mini Kit was used to extract the RNA of each sample following the manufacturer's instructions. Library preparation and Illumina sequencing were performed at the Ecole normale supérieure GenomiqueENS core facility (Paris, France). Messenger (polyA+) RNAs were purified from 250 ng of total RNA using oligo(dT). Libraries were made using the strand specific RNA-Seq library preparation *Stranded mRNA Prep, Ligation kit* (Illumina) and were multiplexed on a P3 flowcell (Illumina). A 118 bp single end read sequencing was performed on a NextSeq 2000 device (Illumina). The analyses were performed using the Eoulsan pipeline (Jourdren et al., 2012), including read filtering, mapping, and alignment filtering. Before mapping, poly N read tails were trimmed, reads ≤40 bases were removed, and reads with quality mean ≤30 were discarded. Reads were then mapped on the *Morpho helenor* genome using STAR version 2.7.8a (Dobin et al., 2013). Alignments from reads matching more than once on the reference genome were removed using Java version of samtools (Li et al., 2009).

<u>Opsin immunochemistry in the eye of *M. helenor* and statistical analyses</u>

Butterflies were euthanized by quickly crushing the thorax and beheading them. In 1x PBS, the heads were bisected and excess tissue was removed under a dissecting scope. The eyes were fixed in 4% paraformaldehyde for 30 minutes in a cold room (approximately 15 °C) with rotation. Fixed eyes then underwent step-wise sucrose baths (10, 20, then 30% sucrose in 1x PBS) for two hours for each step in a 15°C cold room with rotation. The corneal lenses of the eyes were carefully removed under a dissecting scope, and each eye was then embedded in gelatin-albumin blocks. Eyes were oriented so that the slices are presented in distal view. Blocks were then fixed in 4% formalin (in 1x PBS) for 15 hours at 4 °C. Embedded tissue gelatin-albumin blocks were sliced with a Precisionary Compresstome VF-310-0Z to 60 micron slices. Slices were blocked in a blocking solution of 5% (v/v) normal donkey serum and 0.3% triton-X in 1x PBS (0.3% PBST) for one hour at room temperature. The slices were incubated overnight at 4 °C with primary antibodies (1:200 goat anti-BRh, 1:100 guinea pig anti-UVRh, and 1:100 rabbit anti-LW1Rh in blocking solution). Afterwards, the slides underwent five washes with 0.3% PBST for 15 minutes per wash at room temperature. The slides were then incubated overnight in darkness at 4 °C with secondary antibodies (1:250 donkey anti-goat AlexaFluor 488, 1:250 donkey anti-guinea pig AlexaFluor 647, and 1:250 donkey anti-rabbit Cy3 in blocking solution). Again, they were washed five times with 0.3% PBST for 15 minutes per wash and mounted in 70% glycerol. Images were taken using a Zeiss LSM 900 Airyscan 2 confocal microscope under a 20x/0.8NA dry objective in the UC Irvine Optical Core Facility and exported using ZenBlue 3.5. Ommatidium types were then manually tallied.

Raw opsin counts were analysed in R version 4.6.2 using generalized linear mixed-effects models (GLMMs) with a Poisson error distribution. Individual animals IDS were used as a random effect to account for pseudoreplication. Overall differences in composition between sexes were assessed, followed by pairwise comparisons between sexes for each ommatidia and photoreceptor type.

Opsin western blot analyses

To validate the specificity of the custom opsin antibodies, western blot analyses were performed on protein lysates prepared from *M. helenor* (n=20 male and n=16 female) eyes by Boster Bio (Pleasanton, CA). Equal amounts of protein (10 µg per lane) were separated on 4–12% Bis–Tris Plus gels (Thermo Fisher Scientific) by SDS–PAGE and transferred overnight onto nitrocellulose membranes using Tris–Glycine transfer buffer. Membranes were incubated with primary antibodies against UVRh, BRh, and LW1Rh, each diluted 1:1000, for 16–24 h at 2–8°C. Following washes, membranes were incubated with fluorophore-conjugated secondary antibodies, and fluorescent signals were visualized using an Odyssey CLx fluorescent imaging system (LI-COR Biosciences).

PCR amplification protocol of opsin sequences throughout the *Morpho* genus

PCRs were performed on 10 *Morpho* species (*M. amphitryon*, *M. hercules*, *M. aurora*, *M. absoloni*, *M. rhodopteron*, *M. portis*, *M. aega*, *M. zephyritis*, *M. iphitus*, *M. polyphemus*), using DNA samples from collection specimens stored in the National Museum of Natural History in Paris, France. The DNA of those *Morphos* was extracted using the Qiagen extraction DNeasy Blood Tissue Kits. Each PCR reaction was run using 20 µL of solution containing 11.25 µL of water, 4 µL of Buffer 5X for Taq LongAmp, 1.25 µL of dNTPs (6.6 mM), 1 µL of BSA (5mg/mL), 0.625 µL of DMSO, 0.475 µL of primers (10 pM), 0.375 µL of Taq LongAmp (ThermiFischer) and 3 µL DNA. The thermal cycling conditions for the amplification started with a 30s denaturation at 94°C, followed by a gradual decrease of the hybridization temperature during 18 PCR cycles, from 61°C to 52°C, and a 10 min elongation step at 65°C (touch-down method). For the following 50 cycles, the hybridization temperature was fixed to 52°C.

RNA opsin library preparation and Nanopore sequencing methods

To check for opsin RNA expression in different *Morpho* species scattered across the phylogeny of the genus, the eyes of freshly caught males belonging to 5 sympatric species from French Guiana (*M. achilles*, *M. deidamia*, *M. hecuba*, *M. rhetenor* and *M. marcus*), were dissected and stored in RNAlater at -80°C before extraction. RNA was extracted using the Qiagen RNeasy Mini Kit according to the manufacturer's instructions and stored at −80 °C until use. Library preparation and Nanopore sequencing were performed at the Ecole normale supérieure GenomiqueENS core facility (Paris, France). 10 ng of total RNA were amplified and converted to cDNA using a modified version of the protocol described in (Guilcher et al., 2021). Briefly, the ligation and rRNA depletion steps were skipped and the oligo GCAGGGGAAATCATCAGCGTATAACTTTTTTTTTTTTTTTTTTTTTTTTTTTTVN was used for the first strand cDNA synthesis. The oligos TTTCTGTTGGTGCTGATATTGCAAGCAGTGGTATCAACGCAGAGTAC and ACTTGCCTGTCGCTCTATCTTCGCAGGGGAAATCATCAGCGTATAAC were used for the amplification of the full-length cDNAs. Afterwards an average of 15 fmol of amplified cDNA was used for the library preparation. After the PCR adapter ligation, a 0,6X Agencourt Ampure XP beads clean-up was optimised and 2 fmol of the purified product was taken into PCR for amplification and barcodes addition with a 17 minutes elongation at each 18 cycles (barcodes from SQK-PCB111, ONT). Samples were pooled in equimolar quantities to obtain 25 fmol of cDNA and the rapid adapter ligation step was performed. Libraries were multiplexed by 9 on one flowcell FLO-PRO002 according to the manufacturer's protocol. Sequencing was performed with the SQK-PCB111 72-hour sequencing protocol run on the PromethION P2 solo, using the MinKNOW software (versions 5.9.12). A mean of 11 ± 2,8 million passing ONT quality filter reads was obtained for each of the samples. Base-calling from read event data was performed by dorado-0.7.1 sup. Structural annotation was performed following a custom protocol published in protocols.io (DOI: dx.doi.org/10.17504/protocols.io.36wgqd5qyvk5/v1).
