## Supplemental Figures for "Evolution of opsin genes among butterfly species from different forest strata"

### Supplementary Information (Figures): Evolution of opsin genes among butterfly species from different forest strata

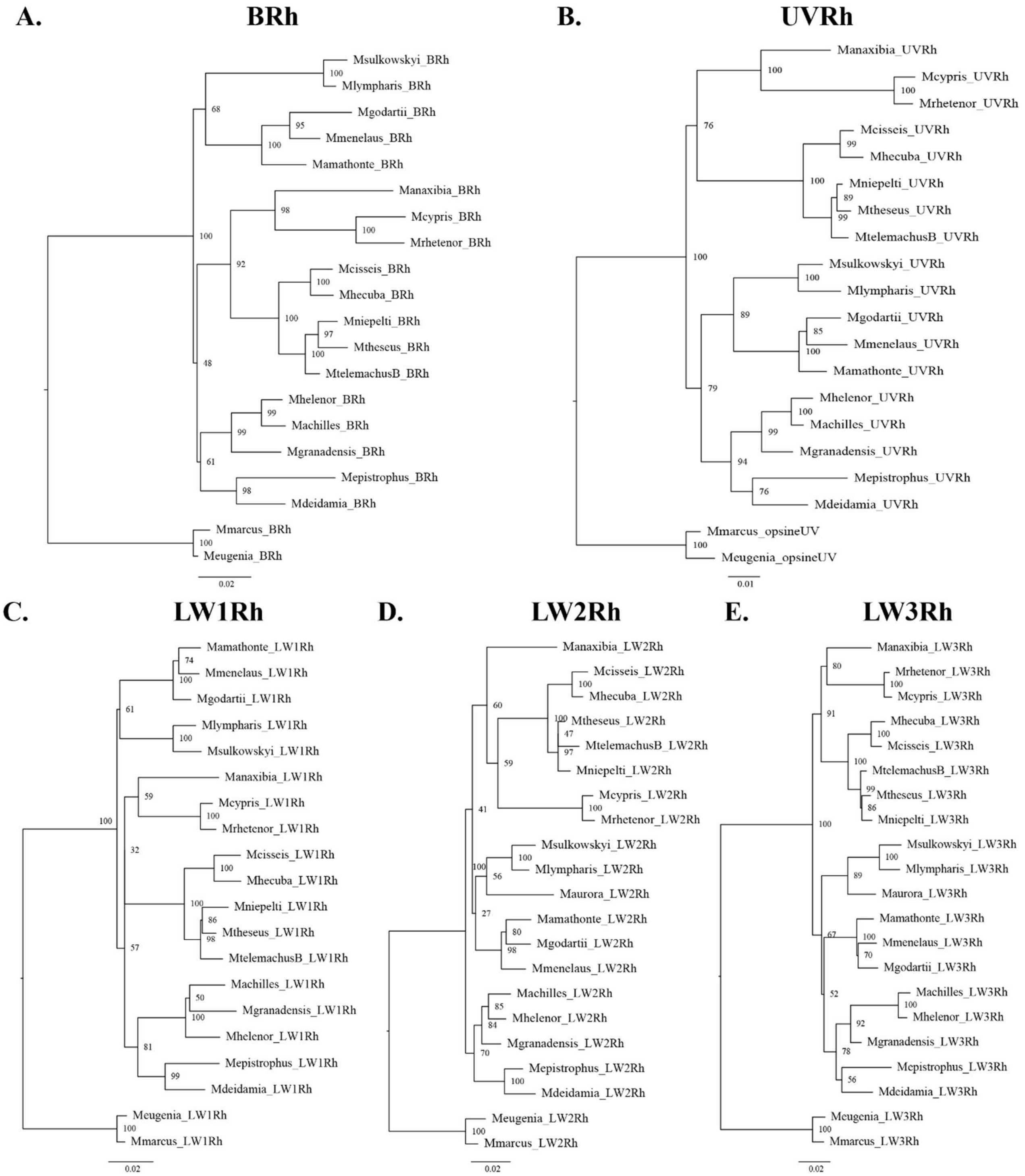


**Figure S1: Gene trees used for the PAML site models.** (A) *BRh* opsin gene tree under the TIM2e+G4 evolutionary model, (B) *UVRh* gene tree under the HKY+F+G4 evolutionary model, (C) *LW1Rh* gene tree under the TIM2+G4 evolutionary model, (D) *LW2Rh* gene tree under the TIM2+R2 evolutionary model, (E) *LW3Rh* gene tree under the TPM3+G4 evolutionary model. The trees presented in this figure are all rooted, but all selection analyses were performed using only unrooted trees.

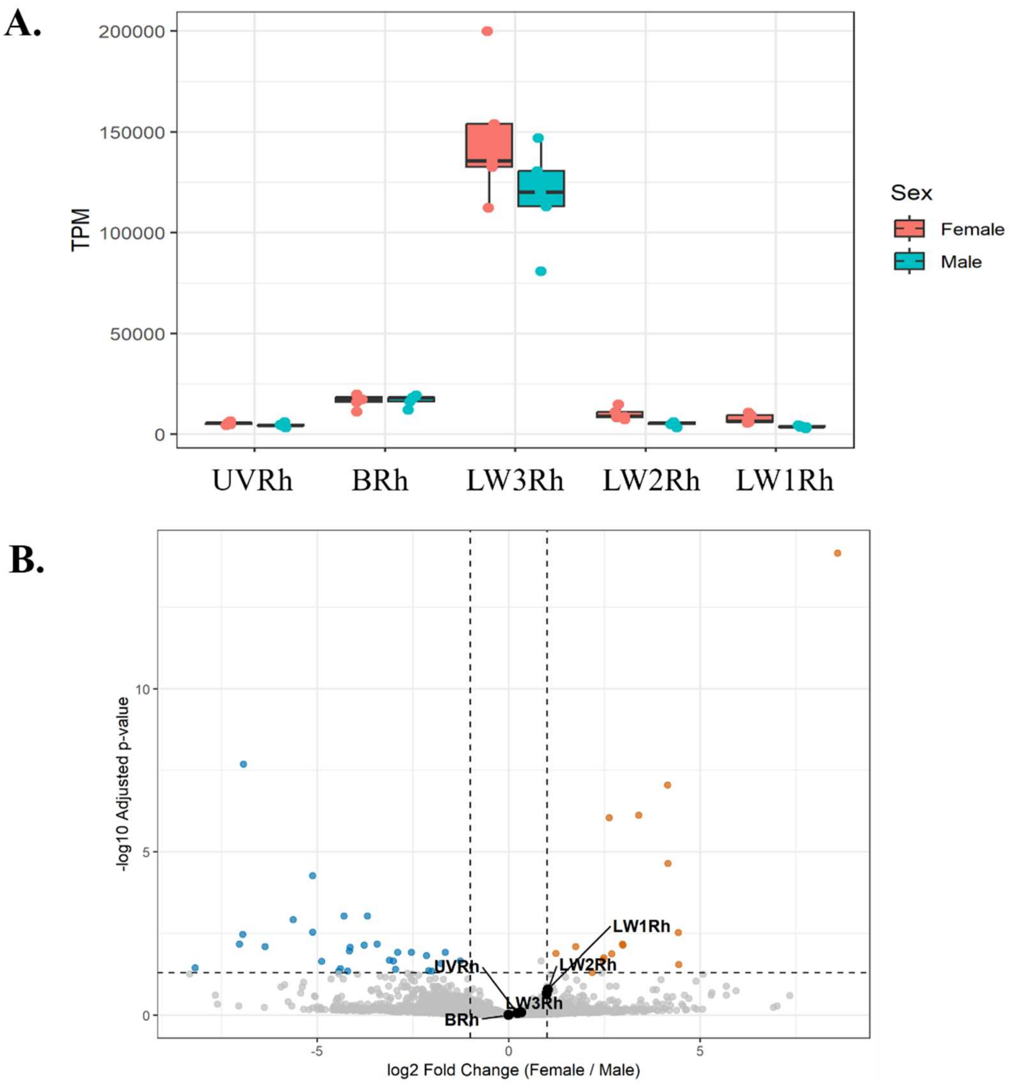


**Figure S2: Level of opsin gene expression in TPM (A) and Volcano plot illustrating the results of the differential expression analysis between males and females**. Orange dots represent the genes upregulated in females, and blue dots the genes upregulated in males. Opsin genes are pictured in black and show no differential expression between sexes.

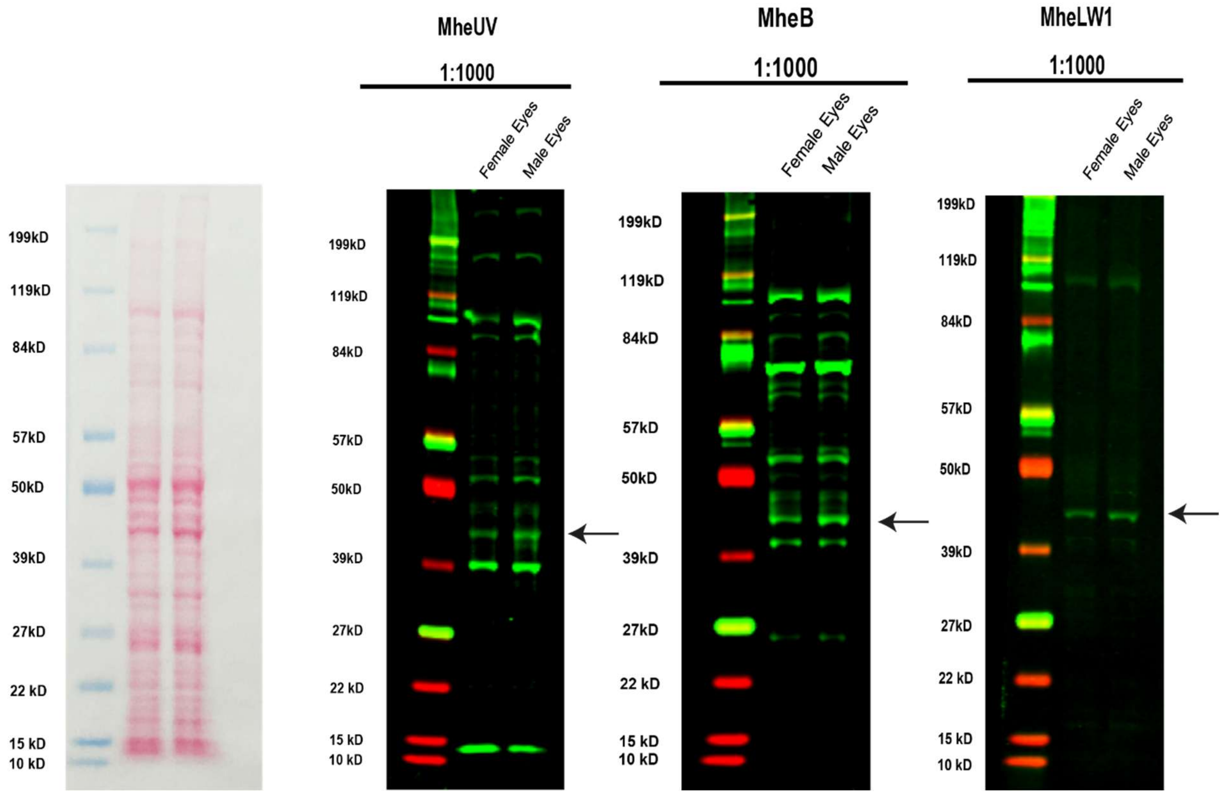


Figure S3: From left to right: General protein stain gel, and guinea pig anti-UVRh, goat anti-BRh, and rabbit anti-LW1Rh opsin polyclonal antibody immunoreactivity on a western blot of *M. helenor* native proteins extracted from adult female (n=16 eyes) and male (n=20) eyes performed by Boster Bio (Pleasonton, CA). An equal amount of protein was loaded in each gel. An immunoreactive band with an approximate molecular weight of 42 kDa, corresponding to the predicted molecular weight of UV, B and LW1 opsins, is indicated by an arrow. Smaller bands might represent alternatively glycosylated sites or proteolytic breakdown products, and larger bands may be opsin dimers or higher order oligomers (Knepp et al. 2012). The peptide used to make the LW1 polyclonal antibody is GSDTGPGIS (see Methods), and the homologous sequences from LW2 and LW3 are SLDPGPGFA and SLDPGPGVA respectively. Given (1) the high dissimilarity in the targeted regions between opsins, (2) the very low amount of dimers observed on the LW1 western blot, and (3) the low brightness of the 42kDa band on the LW1 western blot, there is no evidence of cross-reactivity between LW1 and LW2/LW3.

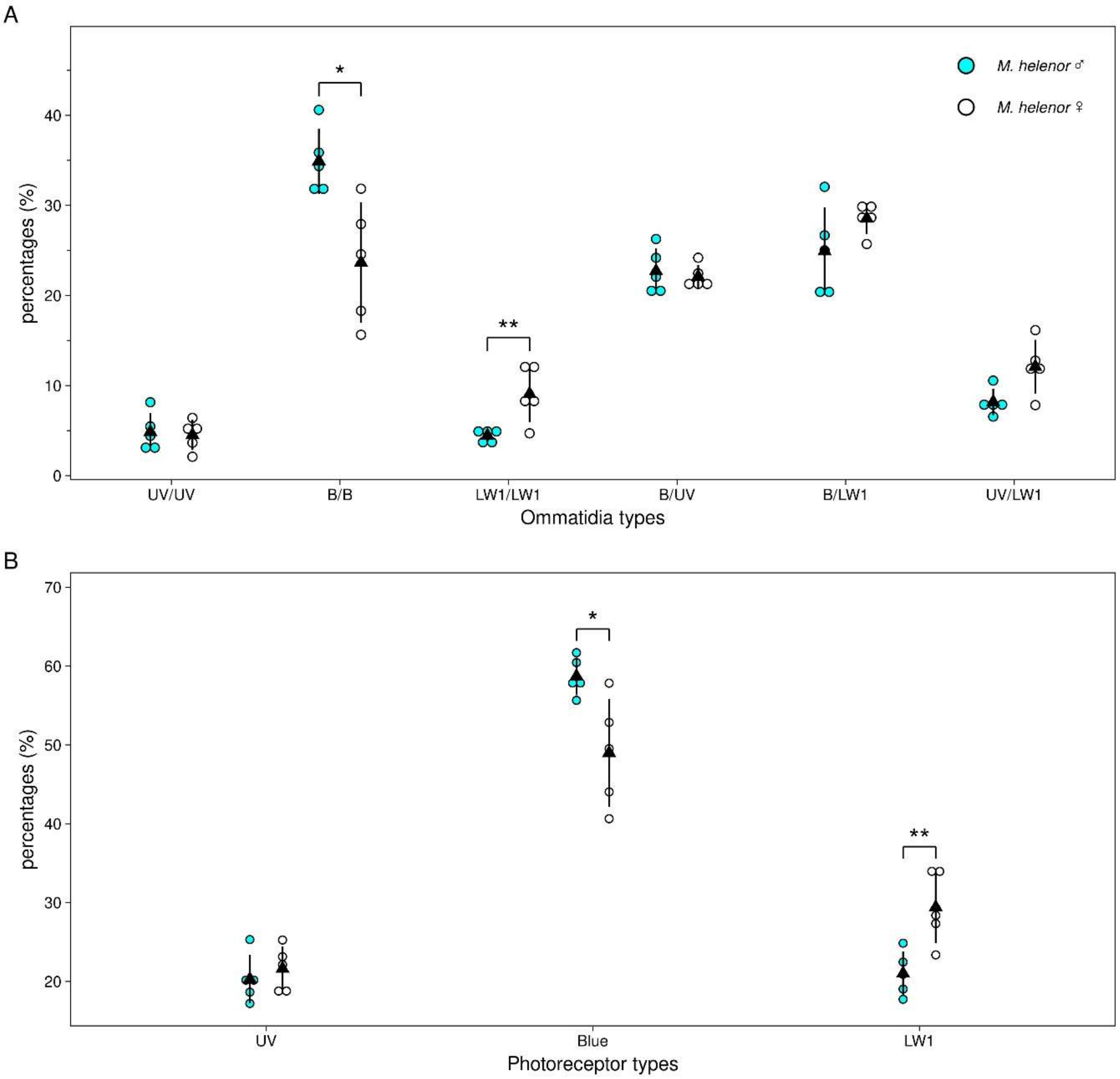


**Figure S4: Frequency of ommatidia (A) and photoreceptor (B) types based on UVRh, BRh, and LW1Rh expression in male (cyan, n=5) and female (white, n=5) *M. helenor* eyes.** Y-axis represents percentages of ommatidia or photoreceptor types in a *M. helenor* eye. Triangles represent mean percentage and error bars represent standard deviations. Sex differences in ommatidia type and photoreceptor type distribution were tested with Poisson generalized linear mixed models with individual animals as a random effect. Distribution differed significantly between males and females for both ommatidia types (sex × type interaction, likelihood ratio test: $\chi^2 = 27.5$, df = 5, $p < 0.001$) and photoreceptor types ($\chi^2 = 17.7$, df = 2, $p < 0.001$). Brackets show female vs. male comparisons within each type, with Holm-adjusted p-values (* $p < 0.05$, ** $p < 0.01$, *** $p < 0.001$).

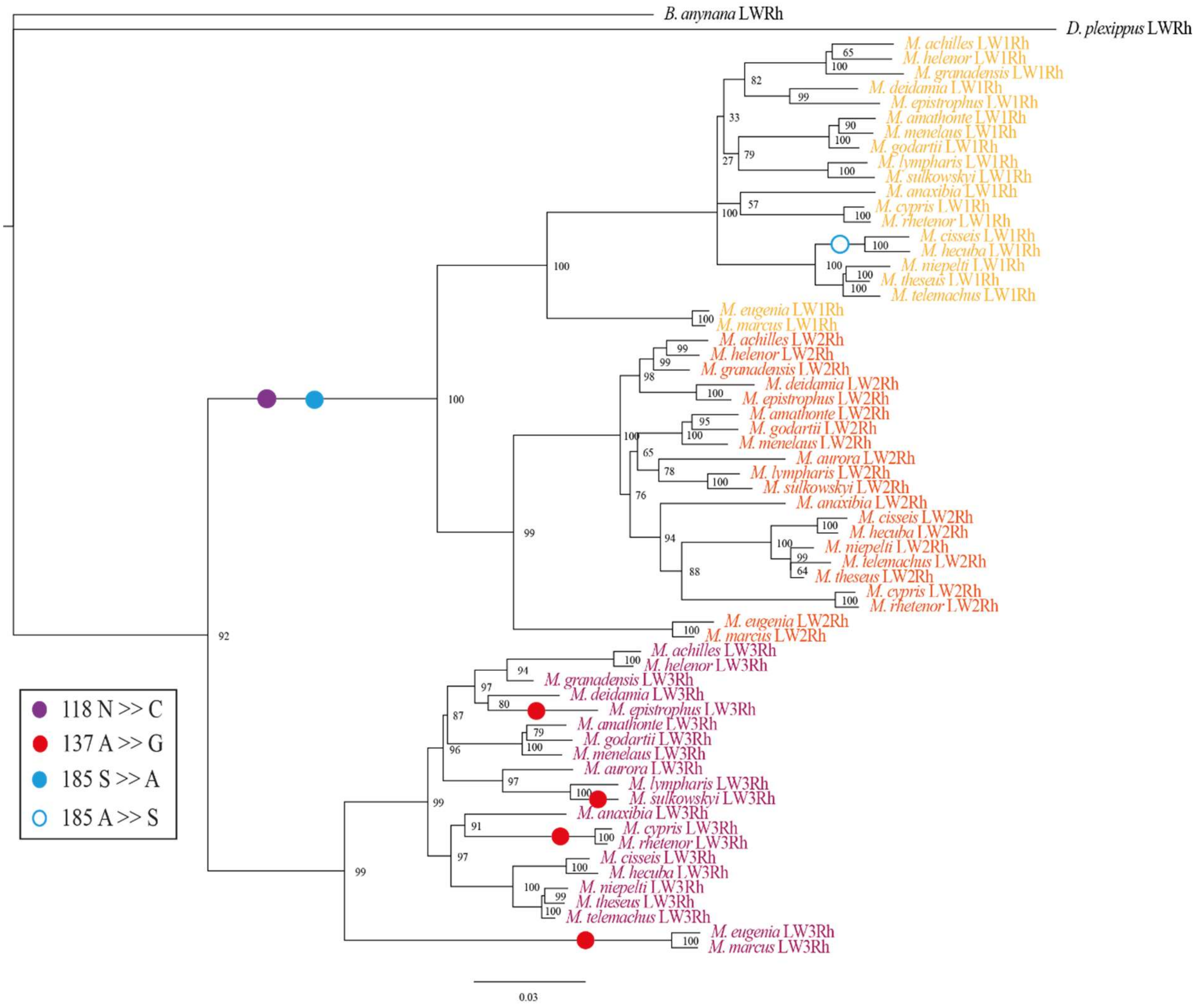


**Figure S5: *LWRh* phylogeny reconstructed using the *LWRh* nucleotide sequences of 21 *Morpho* species, under a GTR+I+G4 evolutionary model.** The long wavelength opsins of *Bicyclus anynana* (GenBank number: AY918895.2, Frentiu et al. 2007) and *Danaus plexippus* (GenBank number: AY605545.1, Sauman et al. 2005) were used as outgroups. *LW3Rh* genes are shown in dark-red, *LW2Rh* genes in orange and *LW1Rh* genes in yellow. Bootstrap values for each node are presented in the tree. We used colored dots to designate amino acid substitutions found on the LWRh sequences of *Morpho* associated with a spectral shift in other butterflies (Frentiu et al. 2007; Saito et al. 2019).

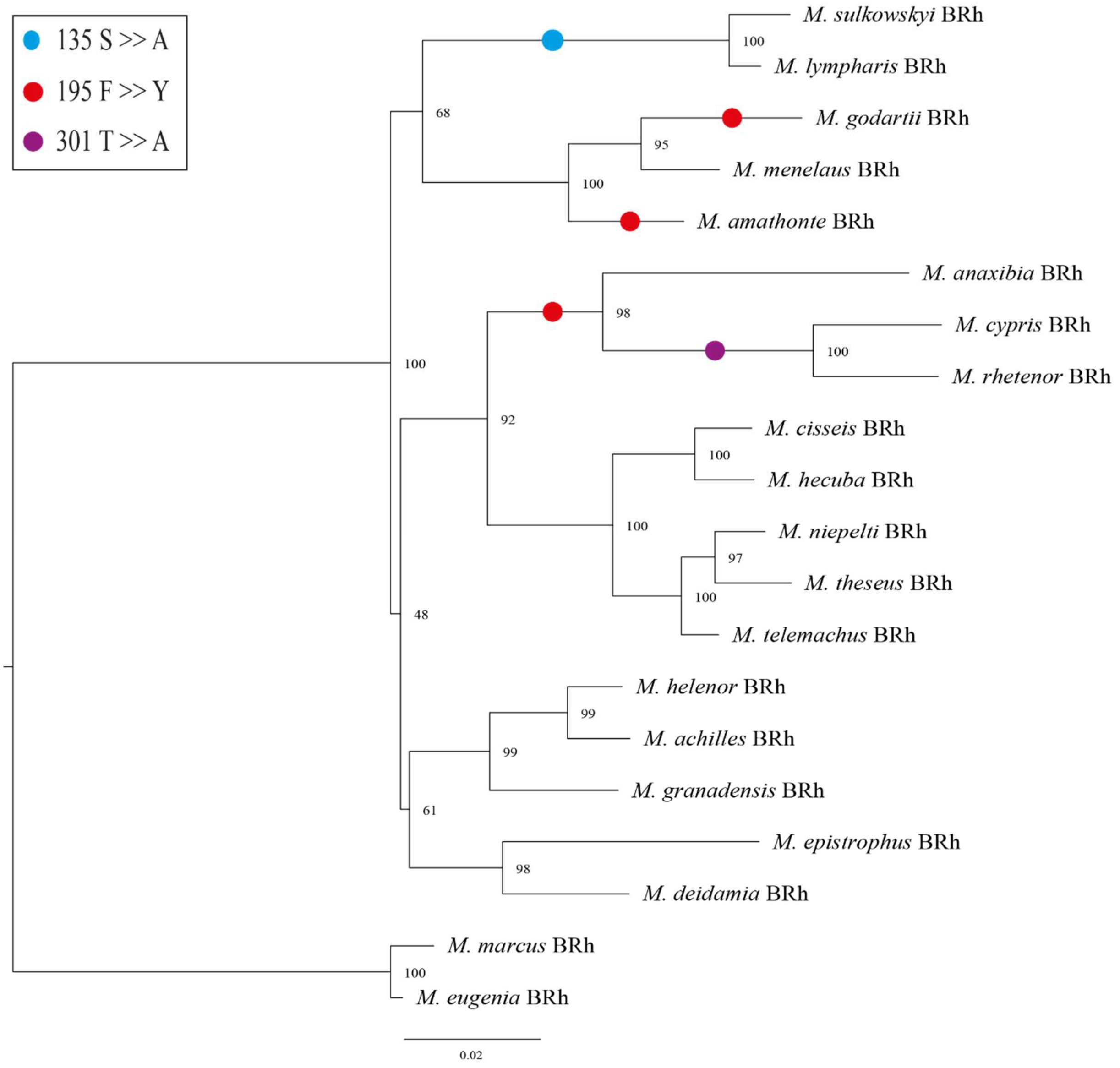


**Figure S6: Amino acid substitutions from the literature identified in *Morpho BRh* genes.** Amino acid positions are numbered according to the *Morpho* sequences but an equivalence table with the numberings from the original sources is provided in Table S5.

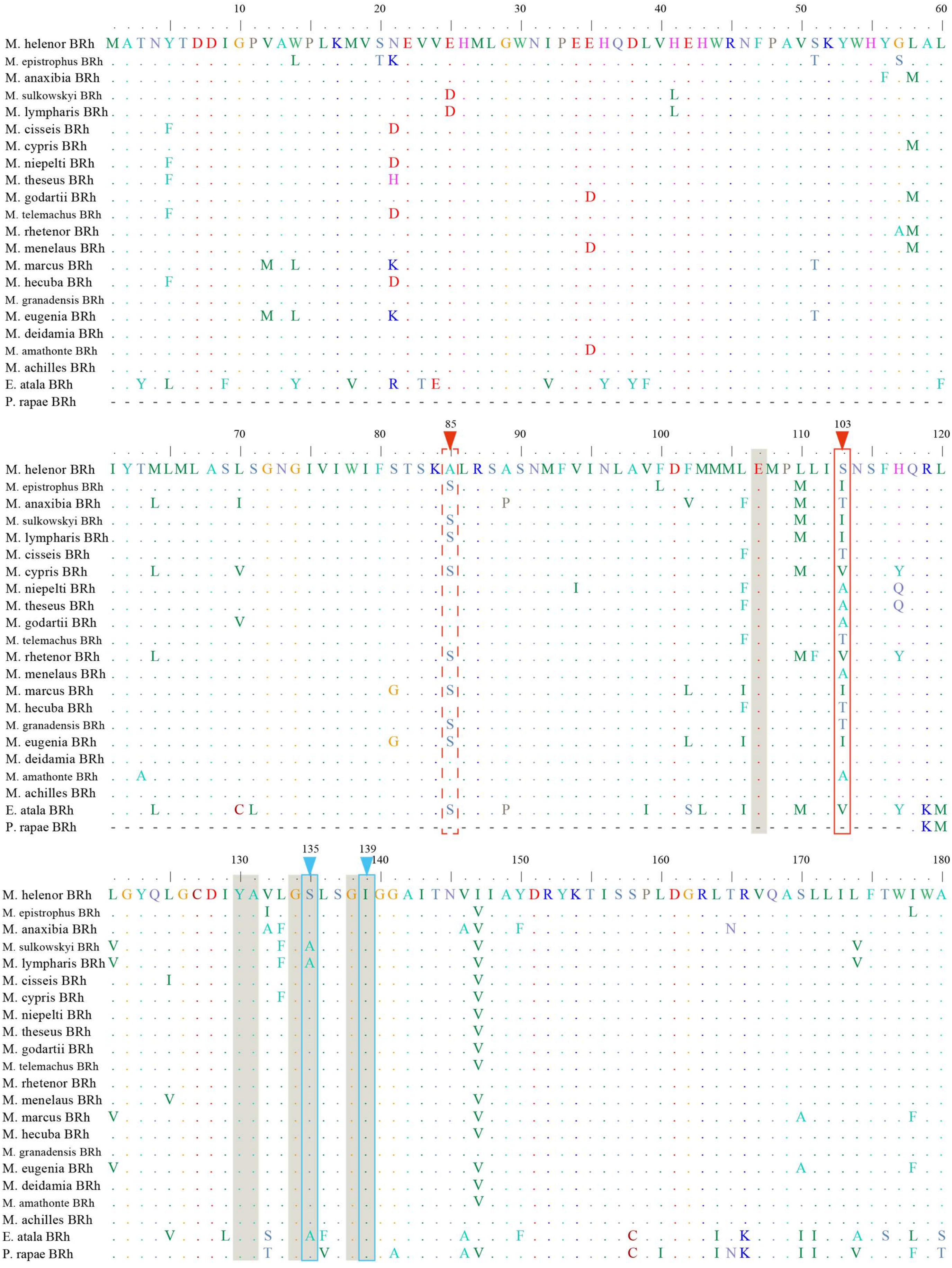
10 20 30 40 50 60
M. helenor BRh MATNYTDDIGPVAWPLKMVSNEVVEHMLGWNIPEEHQDLVHEHWRNFPAVSKYWHYGLAL
85 103
70 80 90 100 110 120
M. helenor BRh IYTMLMLASLSGNGIVIWIFSTSKALRSASNMFVINLAVFDFMMMLEMPLLISNSFHQRL
135 139
130 140 150 160 170 180
M. helenor BRh LGYQLGCDIYAVLGSLSGIGGAITNVIIAYDRYKTISSPLDGRLTRVQASLLILFTWIWA
M. epistrophus BRh
M. anaxibia BRh
M. sulkowskyi BRh
M. lympharis BRh
M. cisseis BRh
M. cypris BRh
M. niepelti BRh
M. theseus BRh
M. godartii BRh
M. telemachus BRh
M. rhetenor BRh
M. menelaus BRh
M. marcus BRh
M. hecuba BRh
M. granadensis BRh
M. eugenia BRh
M. deidamia BRh
M. amathonte BRh
M. achilles BRh
E. atala BRh
P. rapae BRh

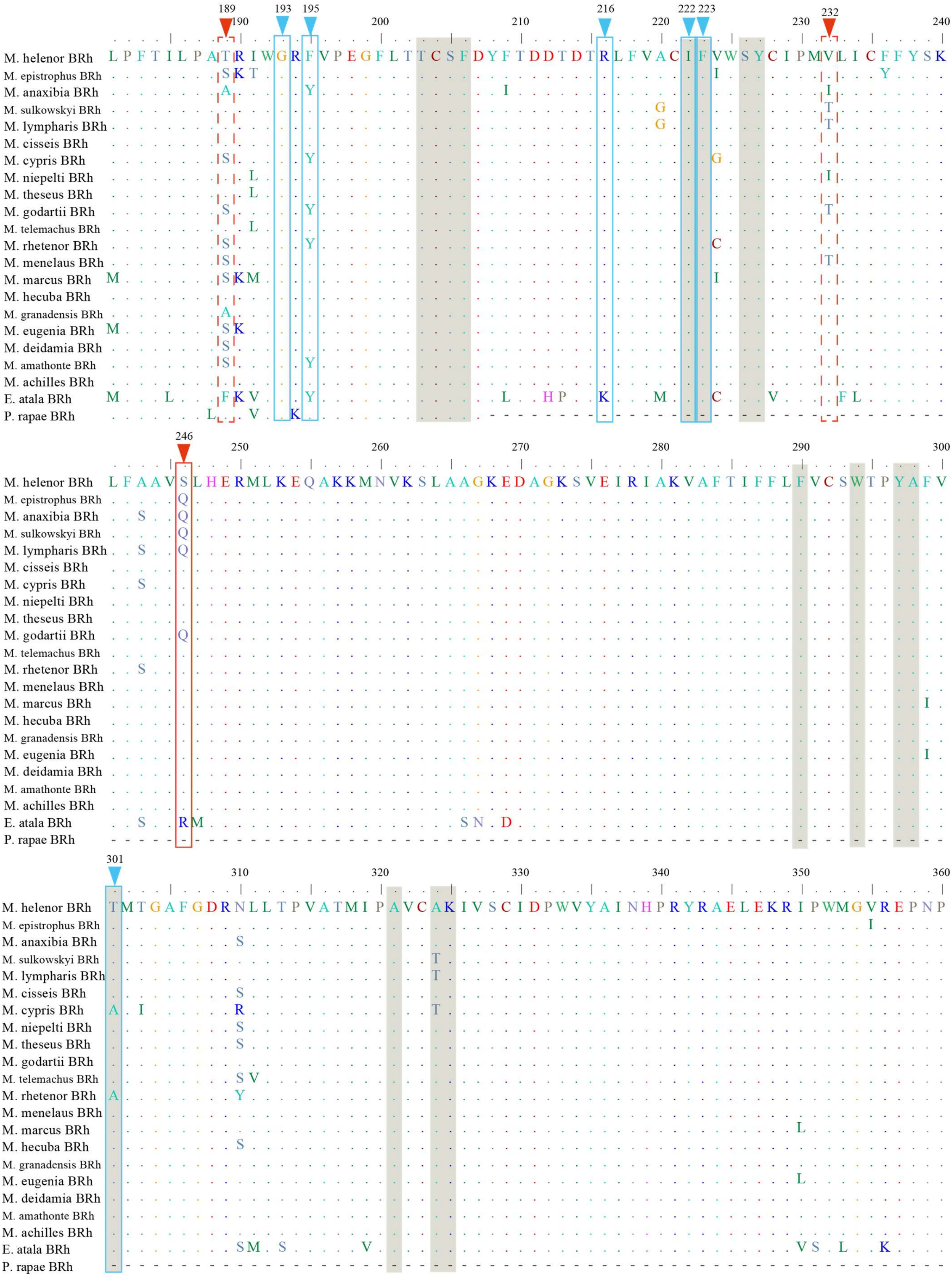

189 193 195 216 222 223 232
190 200 210 220 230 240
M. helenor BRh LPFTILPATRIWGRFVPEGFLTTCSFDYFTDDTDTRLFVACIFVWSYCIPMVLICFFYSK
246
250 260 270 280 290 300
M. helenor BRh LFAAVSLHERMLKEQAKKMNVKSLAAGKEDAGKSVEIRIAKVAFTIFFLFVCSWTPYAFV
301
310 320 330 340 350 360
M. helenor BRh TMTGAFGDRNLLTPVATMIPAVCAKIVSCIDPWVYAINHPRYRAELEKRIPWMGVREPNP
M. epistrophus BRh
M. anaxibia BRh
M. sulkowskyi BRh
M. lympharis BRh
M. cisseis BRh
M. cypris BRh
M. niepelti BRh
M. theseus BRh
M. godartii BRh
M. telemachus BRh
M. rhetenor BRh
M. menelaus BRh
M. marcus BRh
M. hecuba BRh
M. granadensis BRh
M. eugenia BRh
M. deidamia BRh
M. amathonte BRh
M. achilles BRh
E. atala BRh
P. rapae BRh


84

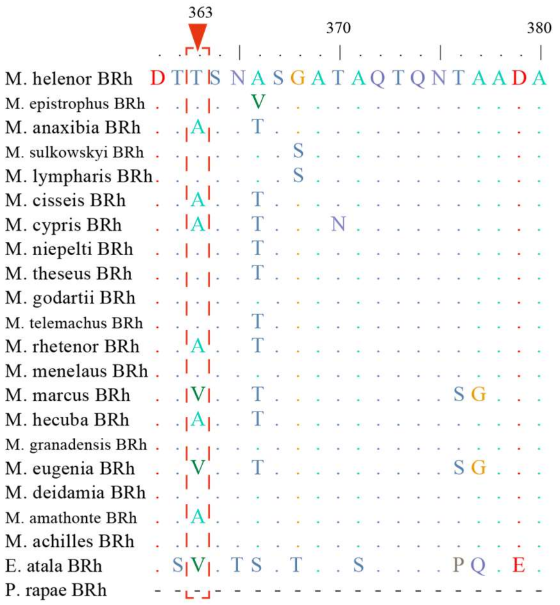


**Figure S7: Alignment of the *Morpho BRh* proteins** highlighting the tuning sites found in the literature (in blue) and sites found to be under positive selection (in red: full lines highlight the AA with a posterior probability > 0.95, and dotted lines the AA with a posterior probability between 0.75-0.95), using *Morpho* AA numbering. The amino acids detected with homology modelling as part of the binding pockets are marked in grey. *Pieris rapae* (Genbank number: BAE19946.1) and *Eumaeus atala* (Genebank number: QPZ55119.1) sequences were aligned to find the AA correspondence and compare our alignment to the tuning site data from Wakakuwa et al. 2010; Frentiu et al. 2015; Liénard et al. 2021.

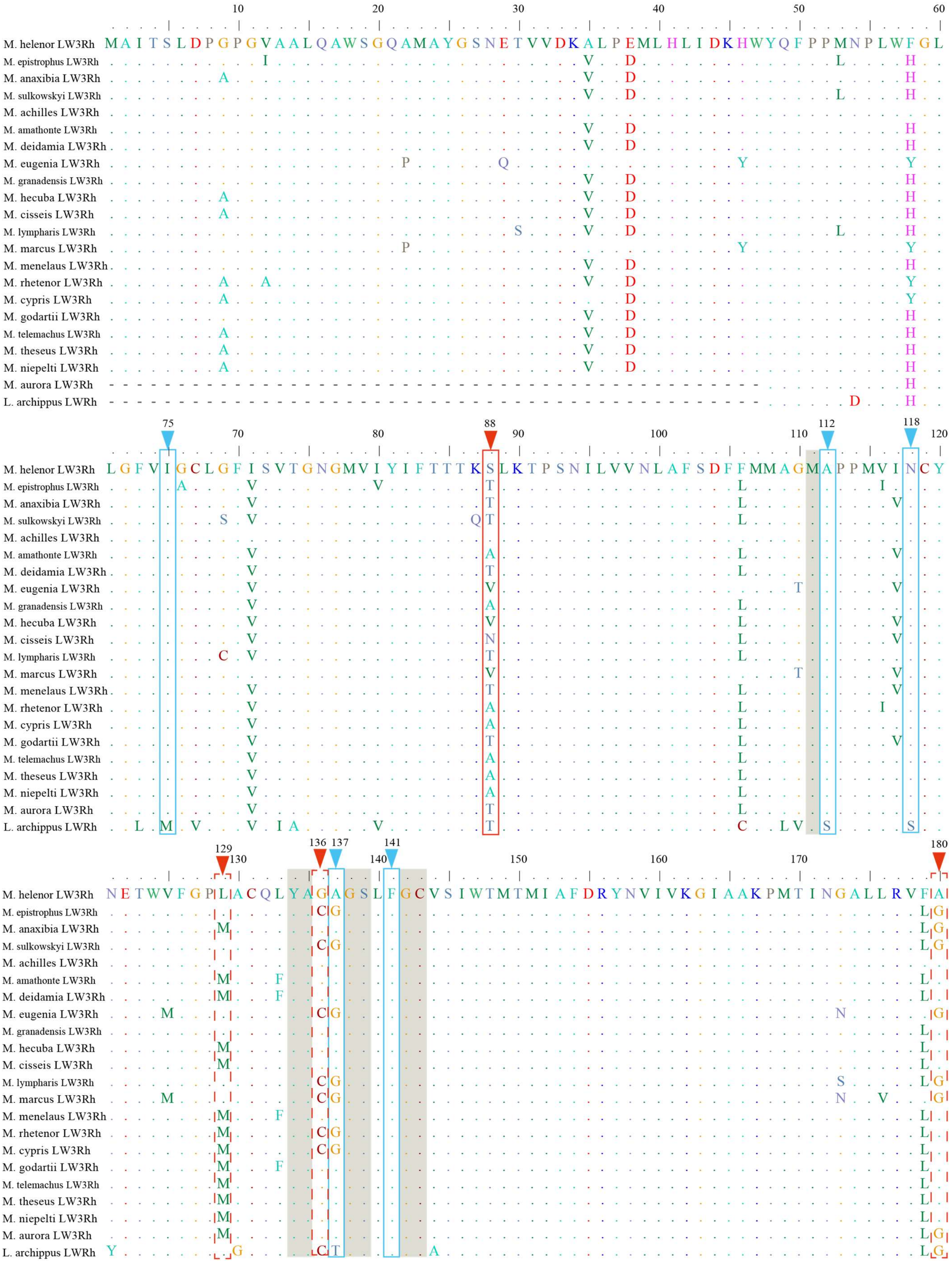
M. helenor LW3Rh
M. epistrophus LW3Rh
M. anaxibia LW3Rh
M. sulkowskyi LW3Rh
M. achilles LW3Rh
M. amathonte LW3Rh
M. deidamia LW3Rh
M. eugenia LW3Rh
M. granadensis LW3Rh
M. hecuba LW3Rh
M. cisseis LW3Rh
M. lympharis LW3Rh
M. marcus LW3Rh
M. menelaus LW3Rh
M. rhetenor LW3Rh
M. cypris LW3Rh
M. godartii LW3Rh
M. telemachus LW3Rh
M. theseus LW3Rh
M. niepelti LW3Rh
M. aurora LW3Rh
L. archippus LWRh
75
88
112
118
129
136
137
141
180

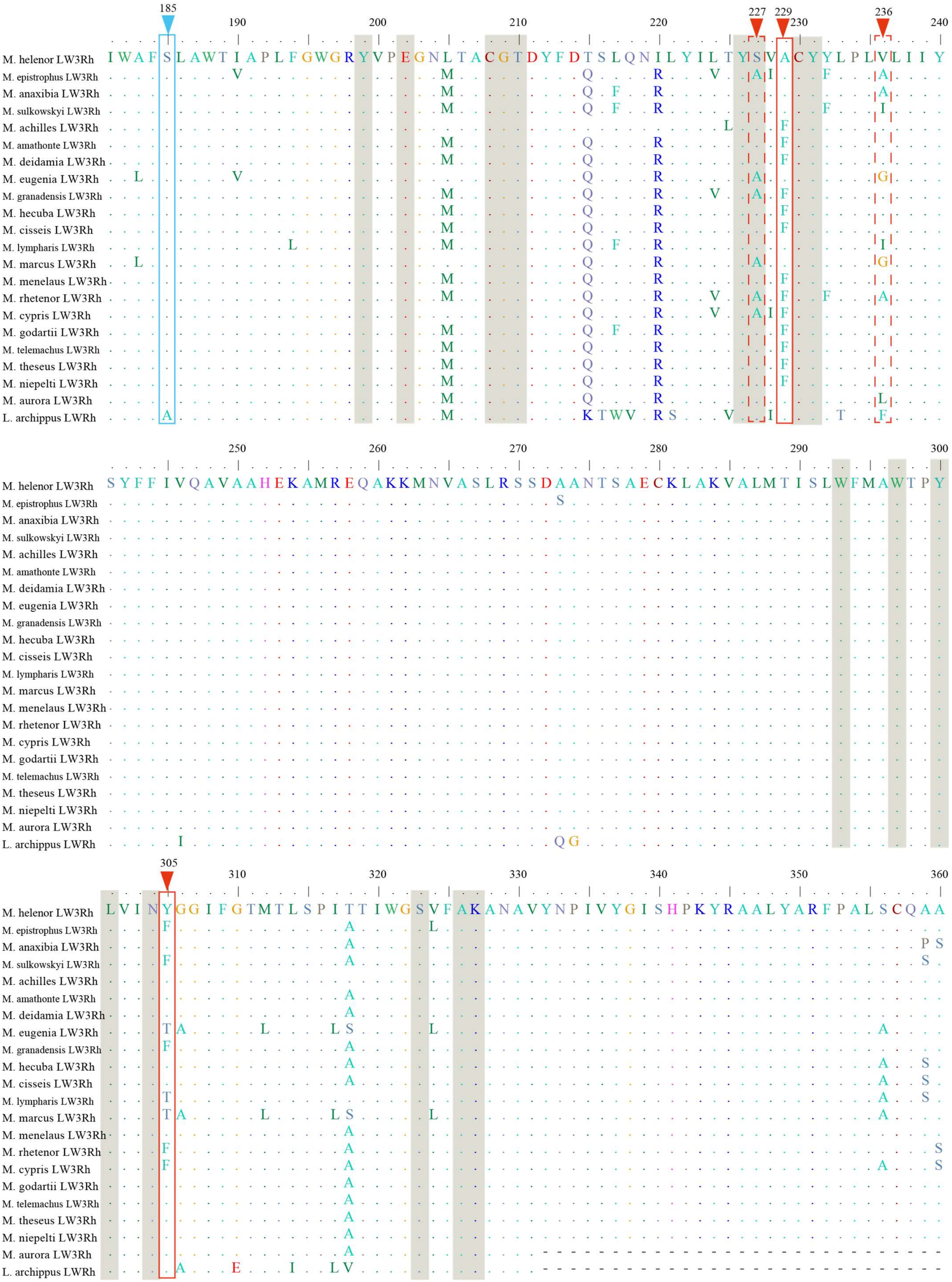
185 190 200 210 220 227 229 230 236 240
M. helenor LW3Rh I W A F S L A W T I A P L F G W G R Y V P E G N L T A C G T D Y F D T S L Q N I L Y I L T Y S V A C Y Y L P L V L I I Y
250 260 270 280 290 300
M. helenor LW3Rh S Y F F I V Q A V A A H E K A M R E Q A K K M N V A S L R S S D A A N T S A E C K L A K V A L M T I S L W F M A W T P Y
305 310 320 330 340 350 360
M. helenor LW3Rh L V I N Y G G I F G T M T L S P I T T I W G S V F A K A N A V Y N P I V Y G I S H P K Y R A A L Y A R F P A L S C Q A A
M. epistrophus LW3Rh
M. anaxibia LW3Rh
M. sulkowskyi LW3Rh
M. achilles LW3Rh
M. amathonte LW3Rh
M. deidamia LW3Rh
M. eugenia LW3Rh
M. granadensis LW3Rh
M. hecuba LW3Rh
M. cisseis LW3Rh
M. lympharis LW3Rh
M. marcus LW3Rh
M. menelaus LW3Rh
M. rhetenor LW3Rh
M. cypris LW3Rh
M. godartii LW3Rh
M. telemachus LW3Rh
M. theseus LW3Rh
M. niepelti LW3Rh
M. aurora LW3Rh
L. archippus LWRh

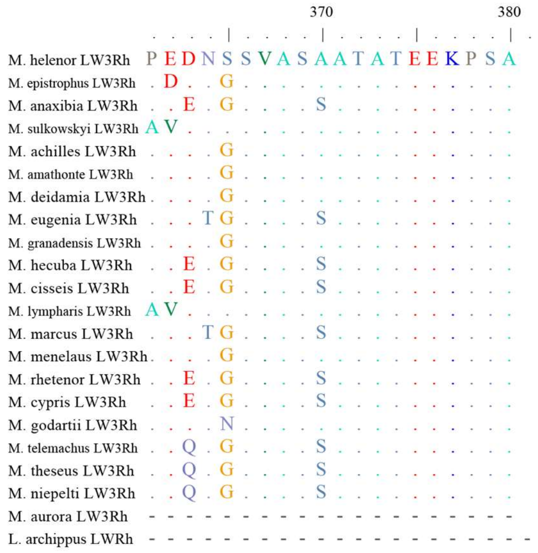


**Figure S8: Alignment of the *Morpho* LW3Rh proteins** highlighting the tuning sites found in the literature (in blue) and site sites found to be under positive selection (in red: full lines highlight the AA with a posterior probability > 0.95, and dotted lines the AA with a posterior probability between 0.75-0.95), using *Morpho* AA numbering. The amino acids detected with homology modelling as part of the binding pockets are marked in grey. *Limenitis archippus LWRh* sequence (Genbank number: ABA44350.1) was aligned to find the AA correspondence and compare our alignment to the tuning site data from Frentiu et al. 2007 and Saito et al. 2019.

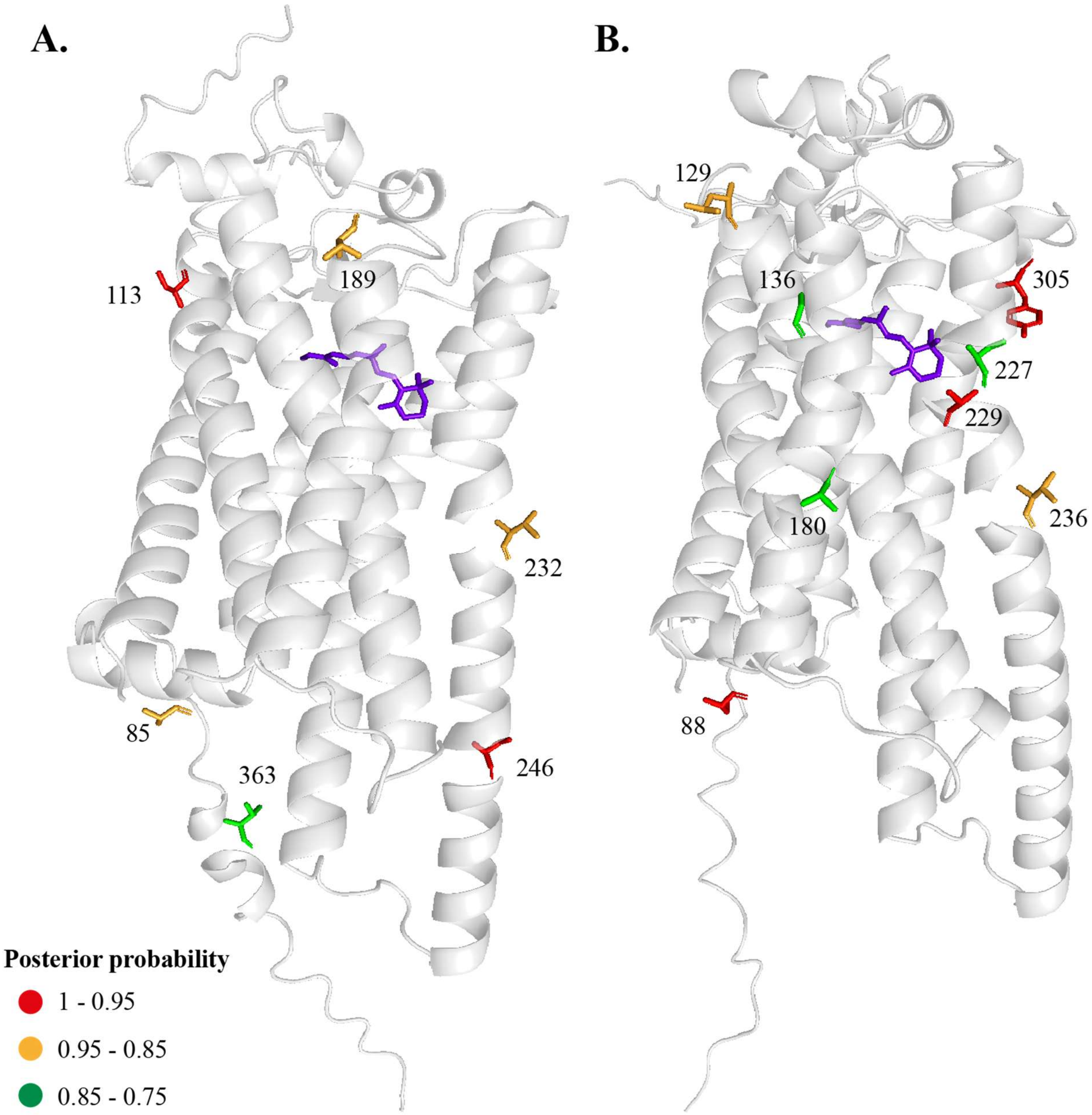


**Figure S9: Positive selection on the BRh (A) and LW3Rh (B) opsin proteins in *Morpho*.** The amino acids detected by BEB PAML's analysis are represented in stick figures, and colored depending on their probability to be under positive selection (posterior probabilities): from 75% to 85% (green), from 85% to 95% (orange) and from 95% to 100% (red). The approximated chromophore location obtained via homology modelling based on the crystal structure of the spider's rhodopsin is pictured in purple.

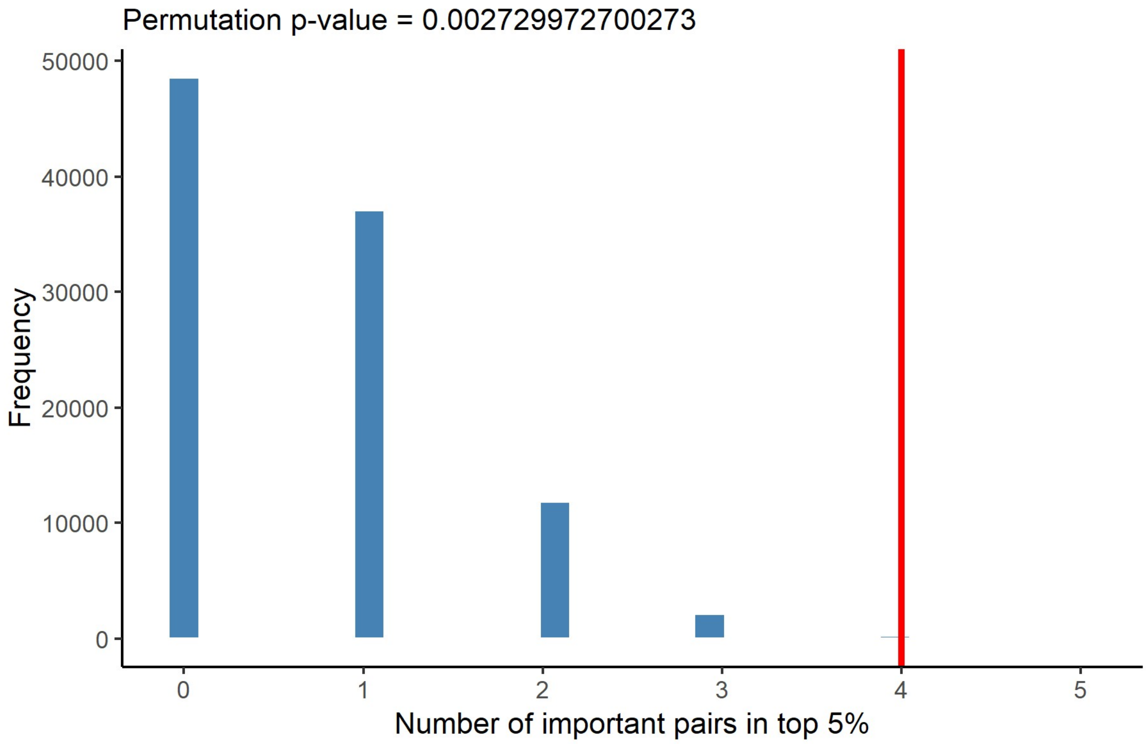


**Figure S10: Permutation test assessing the enrichment of positively selected amino acids among the 5% of amino acid pairs with the smallest p-value detected by EvoScope.** The distribution represents the number of pairs containing positively selected amino acids in the top 5% across 100 000 random permutations. The red line indicates the observed number of pairs with positively selected amino acids. Enrichment was assessed by comparing the observed value with the null distribution generated by permutation. The one-sided permutation p-value is shown.
